# Shoulder muscle recruitment of small amplitude motor units during the delay period encodes a reach movement plan which is sensitive to task context

**DOI:** 10.1101/2022.02.09.479690

**Authors:** Satya Rungta, Aditya Murthy

## Abstract

Understanding how motor plans are transformed into appropriate patterns of muscle activity is a central question in motor control. While muscle activity during the delay period has not been reported using conventional EMG approaches, we isolated motor unit activity using a high-density surface EMG signal from the anterior deltoid muscle to test whether heterogeneity in motor units could reveal early preparatory activity. Consistent with our previous work (Rungta et al., 2021), we observed early selective recruitment of small-amplitude size motor units during the delay period for hand movements like early recruitment of small-amplitude motor units in neck muscles of non-human primates performing delayed saccade tasks. This early activity was spatially specific and increased with time and resembled an accumulation to threshold model that correlated with movement onset time. Such early recruitment of ramping motor units was also observed at the single trial level as well. In contrast, no such recruitment of large amplitude size motor units, called non-rampers, was observed during the delay period. Instead, non-rampers became spatially specific and predicted movement onset time after the delay period. Interestingly, spatially specific delay period activity was only observed for hand movements but was absent for isometric force-driven cursor movements. Nonetheless, muscle activity was correlated with the time it took to initiate movements in both task conditions for non-rampers. Overall, our results reveal a novel heterogeneity in the EMG activity which allows the expression of early motor preparation via small amplitude size motor units which are differentially activated during initiation of movements.

**Significance statement:** We studied the spatial and temporal aspects of response preparation in the anterior deltoid muscle using high-density surface EMG. Our results show that early spatially specific ramping activity that predicted reaction times could be accessed from muscle activity but was absent during isometric force-driven cursor movements. Such ramping activity could be quantified using an accumulator framework across trials, as well as within single trials but was not observed in isometric reach tasks involving cursor movements.

## Introduction

The coupling of neural activity to muscle activity enables the transformation of motor plans into motor execution. To enable tight control over such coupling, the nervous system has evolved multiple possible mechanisms to gate the flow of information between the central nervous system and the peripheral musculature that involves the motor cortex (Kaufman et al. 2014), the basal ganglia (Hikosaka and Wurtz 1983a, 1983b), as well as the brainstem and spinal cord circuitry (Cohen et al. 2010; Prut and Fetz 1999; Selen et al. 2012;). However, despite the presence of gating mechanisms, certain behavioral paradigms that typically evoke reflexive activity have been shown to initiate early recruitment of proximal limb muscles in cats, monkeys, and humans (Fautrelle et al. 2010; Perfiliev et al. 2010; Saijo et al. 2005; Schepens and Drew 2003). These studies also show that stimulus-linked activity can also be observed in shoulder muscles during such rapid reflexive movements (Kearsley et al. 2021; Pruszynski et al. 2010).

In contrast to studies on reflexive movements, early recruitment of muscle activity has not been typically observed for delayed movements (Georgopoulos et al. 1986; Kettner et al. 1988; Lecas et al. 1986; Pruszynski et al. 2010; Requin et al. 1990; Tanji et al. 1988). Nevertheless, delay period activity is observed in motor cortical cells (Churchland et al. 2010; Kurata 1993; Messier and Kalaska 2000; Requin et al. 1990; Shen and Alexander 1997; Tanji and Evarts 1976; Weinrich et al. 1984), as well as in oculomotor areas such as the Frontal Eye Fields (FEF) and the Superior Colliculus (SC) (Dorris and Munoz 1998; Rungta et al. 2021). Such preparatory periods are also correlated with benefits in reaction time (Rosenbaum 1980), raising the possibility that central preparatory activity may have also direct facilitatory effects on agonist muscle activity which may have gone unnoticed in conventional EMG recordings due to their small effects. Consistent with this notion, Mellah et al. 1990 have shown the presence of delay period motor unit activity in axial and proximal skeletal muscles during a delayed movement task in monkeys. More recently, Rungta et al. 2021 have also observed the presence of delay period activity in the neck muscle during a delayed saccade task in head-fixed monkeys, confirming previous reports of anticipatory preparatory activity from surface EMG recordings from the neck splenius capitis muscle of monkeys (Roesch and Olson 2003). Interestingly, in their study, Rungta et al. 2021 observed a heterogeneity of responses amongst motor units, such that only small amplitude sized motor units showed a clear modulation during the delay period. Further, the observed ramping activity was modeled using an accumulate-to-threshold model (Asrress and Carpenter 2001; Carpenter and Williams 1995; Hanes and Schall 1996; Nelson et al. 2016) to provide a link between the observed motor activity and saccade initiation times. Thus, taken together this line of evidence suggests the possibility that small amplitude size motor unit activity may reflect a more direct readout of central activity of motor preparation and planning.

To test these ideas further, we revisited the question of whether the delay period activity can be elicited from arm muscles during reaching movement in humans analogous to what was observed from the neck muscles in monkeys during a delayed saccade task. Motivated by the notion that early muscle activity may be more influenced by smaller motor units that are activated earlier during the preparatory period as a consequence of the size principle (Henneman 1957), we used high-density surface electromyography to assess spatially localized muscle activity that can be used to assess the contribution of smaller motor units that would otherwise be obscured by standard whole muscle EMG activity (Gazzoni et al. 2004, 2004; Merletti and Farina 2009; Rau and Disselhorst-Klug 1997). We also tested whether such delay period activity was sensitive to task context and whether this could explain the absence of muscle recruitment as observed in previous work. In this context, we tested the effects of using an isometric force based task to make cursor movements as no overt EMG activity was observed in such a task despite the presence of delayed activity in the interneurons of the spinal cord (Prut and Fetz 1999).

## Materials and methods

### Subjects

For this study, we recorded from 9 and 11 male subjects (age group: 26-32) for the hand and cursor movement task, respectively. Prior to the experiments, the subjects had to fill the consent form in accordance with the approved protocol of the Institutional Human Ethics Committee, Indian Institute of Science, Bangalore. Subjects carried out their tasks and were paid for their participation in the experiments based on the number of correct trials they performed. Additionally, we present data in supplementary on 5 subjects each using the delayed only task paradigm, for both hand and cursor movements. The results were consistent with this study and have been added to the supplementary section (see link: https://doi.org/10.5281/zenodo.7655975) to strengthen our findings.

### Experimental setup

The two separate setup(s) were used for carrying out experiments for hand and cursor-based movements are described below:

#### Experiment 1: Hand movements

All the experiments in which subjects had to reach out or make an actual hand movement were conducted in a dark room. A sensor along with an LED was strapped on the pointing finger to record hand movements and provide visual feedback. Stimuli were displayed using a 42” Sony Bravia LCD monitor (60 Hz), placed face down on a wooden setup as shown in **<u>Fig S1</u>**. A semi-transparent mirror (25% transmission, 75% reflectance) was placed at an angle below the monitor screen to reflect the stimuli onto an acrylic sheet placed parallel to and below the mirror. The subjects made movements towards the reflected stimuli over the acrylic sheet. The setup was necessary so that the monitor could not distort the electromagnetic field used for detecting hand movements. Subjects could use the chin rest and found no difficulty in carrying out the experiments.

#### Experiment 2: Cursor movements

Subjects carried out an isometric force-driven cursor movement task using a unilateral robotic arm device (*kinarm end-point robot*, *BKIN Technologies Ltd, Canada*) placed in front of them. They sat on the chair with their shoulders and elbow aligned internally with respect to their body in flexed position as shown in **<u>Fig S2</u>** and were instructed to apply force on the transducer at the end of the robotic manipulandum using their shoulders in the absence of an overt arm and hand movement. The manipulandum was set to a locked position at the centre of the device to avoid any movement of the end-point of the robotic arm. Force applied on the transducer (force/torque sensor, with maximum range: 80N; resolution: 0.02N; *Kinarm end point BKIN Technologies Ltd, Canada*) was mapped on to cursor movements which were displayed using a vertical 21’’ LED DELL monitor placed in front of the subjects on top of the experimental device.

### Task and stimuli

For both hand and isometric-force driven cursor experiments, subjects were instructed to carry out visually guided delayed and immediate movement task.

#### Visually-guided delayed and immediate movement task

*Experiment 1:* In the hand movement task, subjects initially explored 16 different locations on the acrylic sheet (workspace grid: see *Fig S1*), while the experimenter calibrated their hand position and monitored their shoulder muscle response using a high-density surface EMG array (for details see section on surface EMG recordings described below). Based on the maximum response of the anterior deltoid muscle, and to maximise the number of trials, two target positions: (i) towards the movement field (90° or 135°) and (ii) away from the movement field (270° or 315°) were mapped onto diagonally opposite locations lying at an eccentricity of 12° from the centre of the screen. To start the trial, subjects had to maintain fixation of the hand only (300±15ms) on a central box (FP: white, 0.6° × 0.6°) until a stimulus (Target: green, 1° × 1°) appeared at the periphery of the screen. No such instructions were provided for eye movements. Subjects had to maintain fixation until the central box disappeared. The disappearance of the fixation box marked the GO cue to make hand movements to the required target location. The time interval between the target and disappearance of the GO cue (delay condition: 1000 ±150 ms and immediate condition: 0 ms), separated ‘where’ from ‘when’ to initiate a movement. After a successful trial, subjects heard a ‘beep’ sound, as a reward, indicating the trial was carried out correctly. An electronic window of ± 2° drawn from the centre of the fixation box and the target were used to assess the online performance. Subjects were rewarded for each correct trial. On average, subjects carried out around 327 ± 15 trials per session (∼ >75 correct trials for each condition: 2 delay periods×2 MFs).

*Experiment 2:* In the isometric force-based cursor movement task, all eight target locations lying at an eccentricity of 12° from the centre of the fixation box were used. Gain values used between force applied on transducer and cursor movements was around ∼2.21 N/cm. The time interval was reduced to 750±150 ms for the delayed condition and was 0 ms for immediate condition. Subjects carried out on average around 302 ± 10 trials (∼>65 correct trials for each condition: 2 delay periods × 2 MFs) for each session.

### Data collection

Experiments were carried out using a TEMPO/VIDEOSYNC system (Reflecting Computing, USA). The codes for generating stimuli were written and controlled using TEMPO. Two separate experimental setup (s) were used for carrying out the experiment. The visual stimuli were displayed using VIDEOSYNC software onto a (i) Sony Bravia LCD (42 inches, 60 Hz refresh rate; 640 × 480 resolution) monitor for hand movements and (ii) onto a Dell LED (21 inches, 60 Hz refresh rate; 640 × 480 resolution) monitor for initiating force-based cursor movements. While subjects were performing the task, we monitored their hand or cursor positions and recorded electromyographic signals from the anterior deltoid shoulder muscles.

### Hand tracking system

An electromagnetic tracker (240 Hz, LIBERTY, Polhemus, USA) was placed on the tip of the pointing finger. The position of the sensor with reference to a source was read out by the system and sent to the TEMPO software in real time. At the beginning of the session, the experimenter would ensure that hand movements were being made to the correct target location and the traces landed within the electronic window (visible only to experimenter), which was centred on the target. The fixation spot and the horizontal and vertical gains of the hand data were fixed as constant values based on the calibration of the system. The average accuracy across different sessions for the hand tracker, while fixating was around 0.0035 ± 0.0001 cm. This was estimated using the jitter while fixating prior to the appearance of the target. Also, the average spatial end point and accuracy of the hand movements to the target locations was 11.32 ± 0.01 cm. Accuracy was estimated by measuring the mean hand end-point location during a 150 ms period of the post hand fixation time across different sessions.

### Mapping force to cursor movements

The manipulandum on the robotic arm was set to a locked position at the centre of the device to avoid any movement of the end-point robot. The force (maximum<30N) applied on the transducer at the end of the robotic arm was mapped onto cursor movements by changing the gains of the analog input (see **<u>Fig S2</u>**) via the Tempo system. For each trial, the colour of the fixation box (colours: red, yellow or green) indicated the force (levels: high, intermediate or low) needed to be applied on the transducer in order to move the cursor from the fixation box to the target in the absence of an overt hand movement (isometric condition).

For setting up the gain values, we calibrated the system and estimated the required force to generate cursor movements by means of vertically hanging weights. A horizontal force was applied onto the manipulandum by hanging known weights from one end using a custom-built pulley-based system. Displacement of the cursor from the centre position was recorded by varying the weights (range: 300 to 2600 gms) for three different gains (values: 0.85, 1.30 and 2.21 N/cm) and used the tempo system to generate appropriate cursor movements. The different force levels that were required to displace the cursor to targets located at 12^0^ eccentricities from centre of the screen by same amount were approximately 10.2N, 15.6N and 26N, respectively (see **<u>Fig S2</u>**). The average accuracy across different sessions for the cursor, while fixating was around 0.19 ± 0.0082 cm. Fixation accuracy was estimated using the jitter while fixating prior to the appearance of the target. Also, the average spatial end point accuracy of the cursor movements to the target locations was 11.91 ± 0.013 cm. End point accuracy was estimated by measuring the mean cursor end-point position during a 150 ms period of the post cursor fixation time across different sessions.

### Hand and cursor onset detection

Hand and cursor onsets were detected offline. The noise level in the position trace profile captured during the fixation time was used as a criterion to set a threshold to mark the beginning of hand and cursor movements. Multiple check points were incorporated in the algorithm to ensure reliable hand and cursor onset detection. First, the cut-off criterion was set at a threshold of 2.5 standard deviations above the average noise level observed in the sensor data during the fixation time for each trial. Second, the onset points from where the hand traces were seen as increasing monotonically for more than 60 ms (15 bins) was marked as the time for the beginning of the hand or cursor movements. Third, trials in which the hand or cursor latency was less than 100 ms or more than 1200 ms were rejected as anticipatory or delayed movements.

### Surface electromyography recordings

EMG activity was measured using a high-density electrode array built by joining three 10 pin Relimate connectors. Out of multiple electrodes, an eight 8-channel array setup (see schematic in **<u>Fig S3</u>**) was used for recording EMG signals. The data was sampled at 10 kHz and band-passed between 10-550 Hz and stored in the Cerebrus data acquisition system (Blackrock Microsystems, USA). The reference electrode was also placed within the array, while a ground electrode was placed on or at the back of the ear lobe of the subject.

### Conduction velocity and extraction of motor units

A high-density array setup (see right panel, **<u>Fig S3</u>**) was used for recording surface electromyographic (sEMG) signals. These signals were analysed in Matlab (Mathworks, R2018b). We estimated the conduction velocity for the surface EMG signals to link spikes observed in the surface EMG signal as potential propagating motor units within the muscle along the electrodes in the array. *Fig S3* shows a snippet of ∼25ms recording showing propagation of motor activity along the electrodes. Signals were captured with respect to a reference electrode located on the same muscle (see *Fig S3*) and analysed in differential or bipolar mode. Using the voltage differences across the electrode locations along the column of the array (DF_1_ = E_2r_-E_1r_; DF_2_ = E_3r_-E_2r;_ DF_3_ = E_4r_-E_3r_), we estimated the conduction velocity for the motor activity. Cross correlation analysis was done on trial-by-trial basis across the session (for an example trial from a representative session, see *Fig S3*) between DF_1_ and DF_2_ to measure the delay between the two signals (lag_1_: 0.8 ms). Conduction velocity was estimated by dividing the average lag for the session with the distance between the corresponding electrode pairs (d_1_=2.5 mm). Similarly, conduction velocity for d_2_=5 mm was estimated from the average lag2 which was measured from a cross correlation between DF_1_ and DF_3_ (lag_2_ = 2.3ms, for the example trial from the representative session). The values for estimated conduction velocity were consistent across the population for different inter-electrode distances (hand: d_1_=2.5 mm: cv_1_ = 1.24±.28 m/s and d_2_ =5.0 mm, cv_2_ = 1.21±.29 m/s and cursor: d_1_=2.5 mm: cv_1_= 1.30±.33 m/s and d_2_=5.0 mm, cv_2_ = 1.21±.20 m/s; see *Fig 3S*).

A threshold criterion was used to extract motor units from the data set. The criterion (2.25 SD) was set based on the initial 500 ms of the data from the beginning of each trial. The procedure was carried out initially using EMGLAB software (McGill et al. 2005). In short, the software collected the template of the waveforms (with window length of 9.8 ms for putative spikes as motor units) and the time stamps for all the local maxima that were above the threshold. Next, principal component analysis (PCA) was used to separate the waveforms based on common and similar features both within and across trials. Generally, eight principal axes were sufficient and was found to be robust to explain more than ninety percent of the variability in waveforms within and across trials for all sessions (**<u>Fig S4</u>**). Initially, a PCA was carried out by pooling spikes per trial and were further clustered into 3 to 4 groups as separate motor units based on feature similarity using a k-means clustering technique. Next, to account for differences and stability in spike waveforms across trials, the average spike waveforms from each cluster were pooled across trials (typically >600; ∼4 per trial) for the given session and further sorted into stable waveforms. PCA and k-means based clustering method was carried out, to isolate (typically, >8 to 15) stable waveforms for each session. Next, using a template matching approach, the minimum least square Euclidean distance between raw spikes and stable units from the previous step were used to sort each raw motor spike into appropriate groups. The isolated units were further pooled based on amplitude size and independent of phase into ∼2 to 3 pooled units (typically, isolated by the experimenter) into large, medium and small amplitude based single-unit motor activity (SUA) for further analyses.

### Exclusion of motor units

Units were discarded if they could not be isolated clearly or if they did not show any modulation or task-related response. Units which were only, bi and tri-phasic were considered for analysis. To remove motion artifacts during sessions, units which were saturated or with amplitude greater than ±5000 µV were not considered for analysis.

### Movement fields

The response or the movement field for the anterior deltoid muscle was defined for all sessions showing movement direction selectivity prior to the onset of a movement (**<u>Fig S5</u>**). For hand experiments, before the session began, subjects were instructed to reach for targets shown in a sequence across the screen in a (4×4) grid. These targets covered the workspace. While the targets were shown, EMG signals recorded from this workspace was used to map the movement field (MF) as described in equation 1 below (see *Fig S1* for representative hand session). Once mapped only two target positions, in and out of the movement field (MF) were shown to subjects to maximise number of trials in a session. The average directional tuning across the population was approximately ∼102^0^ for the preferred direction, indicating a right-upward component, as expected for a shoulder flexor (**<u>Fig S6</u>**). On average, around ∼>75 correct trial per condition (2 delay period × 2 MFs) in each session was used for analysing data in the MF.

For cursor studies, out of the eight target locations, the four positions which showed higher activity during the movement epoch were grouped as inside movement field (in-MF) and the diagonally opposite positions were grouped as outside movement field (out-MF). Since, the MF was large, we combined the results from four different target positions to leverage higher statistical power. The average directional tuning across the population was approximately ∼104^0^ for cursor movement (**<u>Fig S6</u>**). On average around ∼ >65 correct trials for each condition (2 delay period × 2 MFs) were used per session to estimate preferred direction of the MF.

A cosine tuning function was used to identify the preferred direction for each session for hand and cursor experiments. As the MF for the motor units are typically large, target locations which lied within ±90^0^ of the preferred direction was grouped as in-MF. The activity during the movement epoch for all correct trials was regressed with the direction of the target according to equation 1, using the Statistical toolbox in MATLAB.

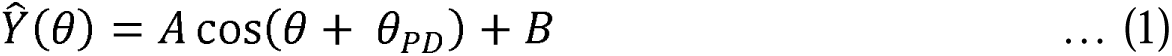

Here, Ŷ(θ) represents the motor activity when the target was presented at an angle θ^0^, B is the baseline firing activity of the shoulder muscles and θ_PD_ is the preferred direction for the motor unit activity.

### Methods for analysing EMG data

We used two methods to analyse EMG data. First, a conventional method such as the root mean square of the EMG signal was used to analyse the recruitment of anterior deltoid (shoulder) muscles. Mathematically, the root mean square measures the variability present in the signal and is thought to reflect information about the recruitment of motor units. A running window of length 35 ms with steps of 1 ms was convolved with the square of the data. The square root of the convolved data for each trial was then used for further analysis.

Second, additionally, we isolated spike waveforms from the EMG signal and extracted their time of occurrence and constructed a spike density function (SDF) from this data. The mean activity pattern of the motor units could be estimated by averaging the convolved data over multiple trials. The spike trains were convolved in both the forward and backward directions using a gaussian filter with a std. deviation of 10 ms. One advantage of processing signals using the above method is that all the features in the filtered data are exactly where they occurred in the unfiltered format. Henceforth, in the results the SDF is referred to as motor activity. Both these approaches (RMS and SDF) led to comparable results (**Fig 1**; Pearson’s r: 0.73; p<0.001).

**Fig 1:**
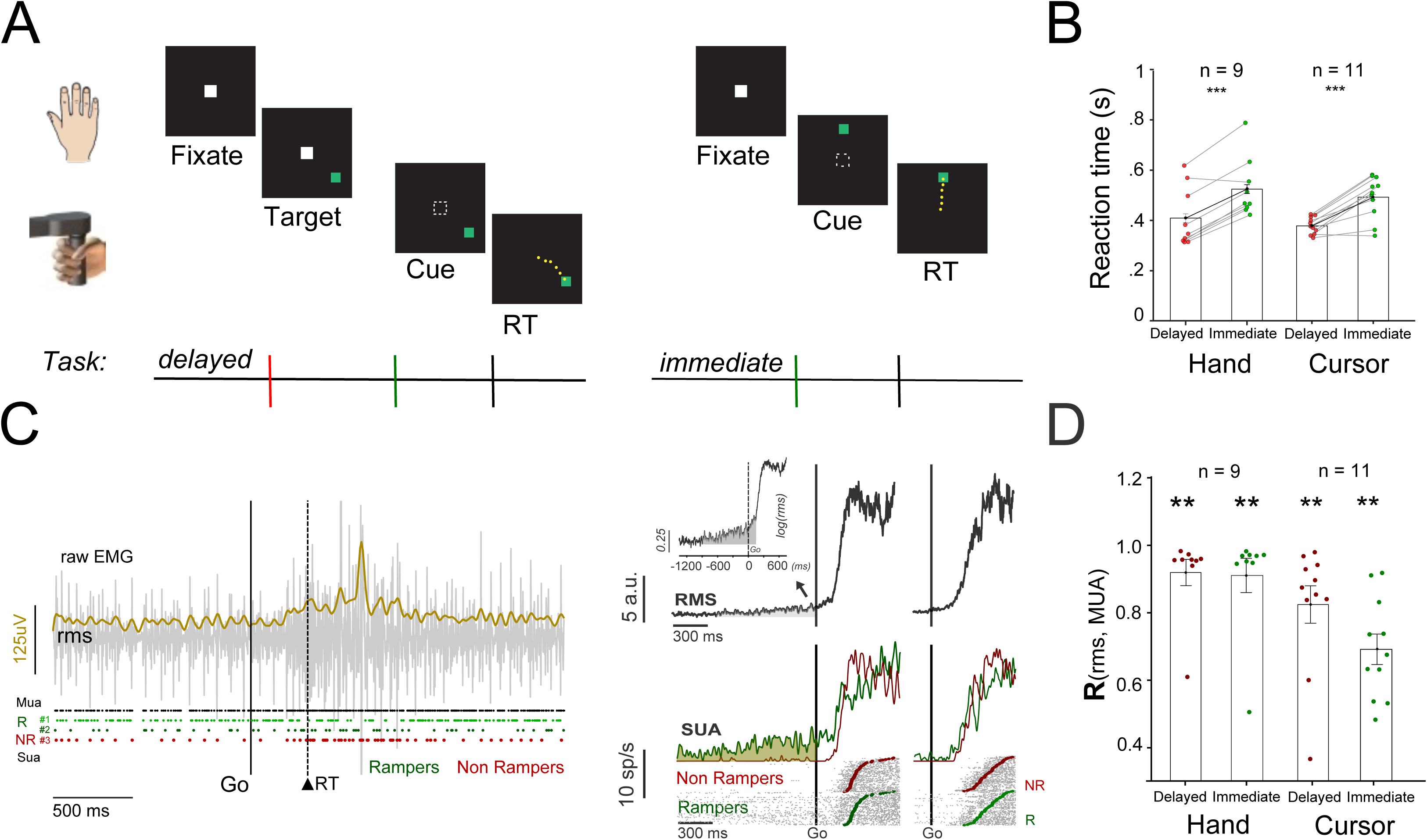
Task and EMG activity. A) Hand and Cursor movements: Subjects made delayed or immediate movements to the green target after the disappearance of the fixation box (GO cue) to initiate a movement. The epoch between the red and green lines demarcates the hold time or the delay period, whereas the epoch between the green and black lines marked the reaction time period. B) Bar plots show the average reaction time to initiate delayed and immediate movements for the hand and the cursor task, respectively. C) *Left panel* The raw surface electromyographic (sEMG) signal for a representative trial recorded using a high-density surface array in differential mode. Dashed lines in black mark the reaction time following the GO cue for the respective trial. The solid trace in yellow shows the root mean square (RMS) variability present in the EMG signal. Black dots mark the onset time for all the motor spikes observed in the EMG trace. Green and red dots show the spike train for spikes separated based on their waveforms. *Right panel*. Shows the average muscle activity for the representative session, aligned to the time of the GO cue analysed using (a) a conventional root mean square method (top, RMS) and (b) a raster-based approach (bottom, pooled single unit activity or SUA separated by amplitude). The inset at the top shows the root mean square activity on a log scale. In the bottom panel: red and green represent the activity for different type of motor responses captured by pooling large and small amplitude size motor units during the session, respectively. Gray marks the time of occurrence for individual spikes. Filled green and red circles represent the time for initiation of movements following the GO cue for rampers and non-rampers, respectively. F) Bar plots show the Pearson’s correlation coefficient between the root mean square of the EMG signal and multi-unit motor activity across all sessions for both hand and cursor tasks. (Correlation between the top and bottom rows for right panel in *Fig 1C* across all sessions).

Since the captured spike density function (SDF) was sparse and had low firing rates, we focussed our analysis on revealing the gradually accumulating activity, independent of underlying oscillatory activity, we used a standard ‘Additive time series decomposition model’ to split the motor activity into its underlying components, namely - trend, cycles and residuals.

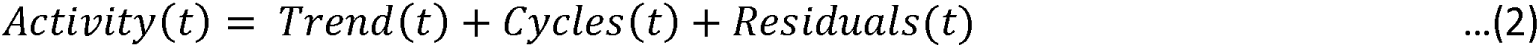

A moving average using a window of 15 bins or 15 ms was used to estimate the trend component present in the time series data. A maximum of 2 to 3 peak cyclic components were identified as cyclical patterns and the remaining irregularity components were verified to be residuals or as white noise after using an auto regressive and moving average (ARMA) approach.

### Direction discrimination time (DDT)

A standard receiver operator characteristic (ROC) analysis (Corneil et al. 2004; Pruszynski et al. 2010; Thompson et al. 1996) was performed on the SDF or motor activity for each time bin after the response (MF-in and MF-out) were aligned on the GO cue. The area under an ROC curve ranges from 0.5 to 1. For each time bin, an ROC value of 0.5 would imply that the motor activity could not be reliably distinguished between in and out of the movement field, whereas an ROC value of 1 would mean complete separation between the two distributions. A threshold criterion during reaction time was set at an ROC of 0.7. The time at which the ROC value crossed the threshold for a continuous period of 150 ms was marked as the direction discrimination time (DDT). Since the ROC time series had cyclic components, it was smoothened using a uniform filter of 45 ms with zero phase shift and then minima between a period of 850 ms prior to GO cue and 25 ms after GO cue was chosen as the starting point to demarcate when the ROC trend increased monotonically during the delay period.

### Categorisation of trials based on different parameters

Reaction time was used as the metric for assessing the initiation of movements. For each session, the data was divided into 2 different groups. Based on this division, different trials were categorised into different conditions - slow and fast reaction times (RTs). Trials with a RT less than 45th percentile were classified as fast RT trials, whereas trials above the 55th percentile onwards were grouped into slow RT trials.

### Components of the accumulator model

Next, we wanted to test whether a linear accumulator model (Carpenter and Williams 1995; Hanes and Schall 1996; Noorani and Carpenter 2016; Reddi and Carpenter 2000) could be extended to the periphery to study the initiation of movements. We chose a particular form of accumulation called the LATER model (Linear Approach to Threshold with Ergodic Rate). The LATER model assumes that the appearance of a stimulus triggers a neuronal preparatory signal to rise linearly until it reaches a threshold value, when a response is initiated. The model has also gained some support from neurophysiological studies of saccade eye movements, where the activity of movement neurons in the frontal eye fields (FEF) of monkeys rises steadily prior to a saccade with a rate that varies randomly from trial to trial and predicts the ensuing latency, whereas the final level of activation is relatively fixed regardless of RT (Hanes and Schall 1996; Rungta et al. 2021).

Four important components of this framework that determine movement initiation times are the activity at baseline, the onset of accumulation, the growth rate and the threshold, which were measured for each motor units based on their SDF, under MF-in conditions for - fast and slow RTs. For both hand and cursor movements, a fixed interval of 125ms was used to estimate *the baseline activity (−100 to 25ms), activity at GO cue (−100 to 25ms) and the threshold activity (−150 to -25ms)* after aligning the spike density function on the onset of target, the GO cue and movement onset, respectively. Slopes and onsets were calculated using a regression-based method to estimate the growth rate and the mark the onset or start time for change in the spiking activity (see **<u>Fig S13</u>**). A fixed interval between 75ms following GO cue and until the mean RT for each individual session was used as an initial seed for approximating the slope and onset. The results obtained using the initial regression were interpolated to calculate the time at which the initial slope intersected the spike density function. The global minima from the intersect time was used as the time to mark the onset for change in the spiking activity. In the final step, the slope of the regression line was used, from the marked onset time until the peak spiking activity within the average RT, to estimate the growth rate for the spiking activity. A similar approach was used to estimate slopes and onset for the spike density function prior to the GO cue and movement onset. A fixed interval of 550ms and 250ms was used from 25ms up to 575ms and 25ms up to 275ms prior to the GO cue and movement onset, respectively, to analyse ramping activity during the delay period and prior to movement onset.

### Poisson spike train analysis for single trial onset detection

We carried out analysis at the single trial level to analyse the onset time for spiking activity using a modified version of the Poisson spike train method described previously by Hanes et al. 1995; Legéndy and Salcman 1985. Firstly, we observed that the spike count for different epochs came from a Poisson distribution with different means, next we observed that the distribution for the inter-spike interval (ISI) were significantly different during the periods before and after the GO cue and it could be approximated using an exponential fit. Both these conditions indicated that the spike train during the intervals followed a Poisson process (see **<u>Fig S9</u>**). We ran each trial through the onset detection algorithm previously used and described by Hanes et al. 1995, to mark the probable onset times for ramping or burst activity before and after the GO cue, respectively. Fixed intervals of 550ms prior to GO cue and 50ms after GO cue and 75ms prior to GO cue till 750ms after the GO cue were used as boundary conditions for analysing changes in spiking activity during the delay and RT periods, respectively.

As shown in **<u>Fig S10</u>** and assuming inter-spike interval to follow a Poisson process, the improbability or rather the probability for finding more number of spikes than actually observed was calculated using the mean expected spike count for each spike time interval(s) during the epoch (equation 3). Next, the calculated improbability was log transformed and negative of the value was used as a metric to measure the surprise index (SI) for the spike train as shown in equation 4. A larger SI indicated a low probability that the increase in spiking activity was a chance occurrence. The following steps were next carried out to optimise the SI and to mark the probable onset and end time for the ramping spike activity. First, the spikes were indexed in increasing order of their occurrence in the interval and the SI was calculated for each spike in the spike train. The spike having the maximum SI was marked as the end of burst or end of the ramping activity. The same process was next carried out from the end of ramping action but now in reverse order (end of the ramping activity or the last spike was indexed as the first spike) and SI was calculated for each spike from last to first in decreasing order. The spike which had the maximum SI was marked as the beginning of the ramp. To be more conservative in the approach, the algorithm continued indexing the spikes further till the SI value dropped below the significance level to mark the onset or beginning of the ramping activity for the given spike train. The equations used for calculating the probability and SI are:

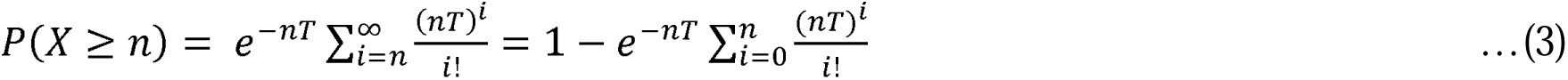

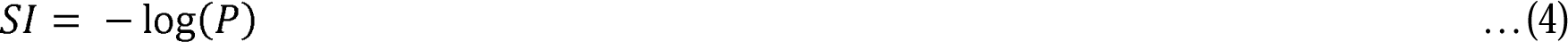

### Statistical tests

All statistical tests were done on population data across subjects (unless specified). A Lilliefors test was carried out for checking normality of the data, under each condition. A N-way ANOVA was carried out, when necessary, where n represents the number of multiple factors. A standard t-test or paired t-test was carried out whenever required. If normality failed, a sign test was done instead of paired sample t-test, and a Kruskal Wallis test was done for ANOVA. A Binomial test was done to test whether the trend in the data was above chance level. In-built MATLAB functions and the statistical toolbox were used to carry out appropriate analysis.

## RESULTS

Subjects performed visually guided delayed and immediate hand or cursor movements (**Fig 1A**). An eight (8) channel high-density electrode array was used for recording surface electromyographic (EMG) signals from proximal shoulder muscles (anterior deltoid) while the subjects performed the task (see methods for details). As known from previous findings, we observed that, the delay period, prior to the GO cue in the delay time condition, was used by the subjects to prepare for an upcoming movement. Significantly shorter reaction times (RTs) (**Fig 1B**) were observed for delayed movements when compared to immediate movements (delayed task: 397.54±36.63ms and immediate: 511.13±35.24ms; t(8) = -7.28, p<0.001). These results indicate that some aspect of motor preparation occurred during the delay period which conferred a reaction time advantage.

### *Experiment 1*: Hand Movements

We analysed the EMG activity through two distinct methods: (a) the traditional root-mean square approach and (b) the raster-based approach. In the former approach, a running window was used to capture the root mean square variability as the envelope of the EMG signal (see methods, top right panel: **Fig 1C**), whereas in the latter approach, EMG signals were first decomposed into motor spikes and separated into multi-unit and pooled single motor activity based on amplitude size. Spike timings were convolved using a temporal gaussian filter for obtaining a spike density function (see methods, bottom right panel: **Fig 1C**). Both approaches gave comparable results (**Fig 1D**; delayed task: Pearson’s r = 0.90± 0.04, p<0.0001 and immediate: Pearson’s r = 0.90 ± 0.02, p<0.0001). Since the latter approach, provided additional information about the spike waveforms and the frequency of spike occurrence, we used the raster-based approach henceforth to analyse our results, unless otherwise specified.

### Recruitment of motor units based on amplitude size and baseline activity

We typically isolated 2 to 3 pooled motor units (maximum of 3) from surface EMG recordings analysed in differential mode from one of the channel combinations available in the array per subject. As shown for the representative session in **Fig 2A**, we isolated different amplitude size motor units. These motor units showed two different kinds of motor responses: rampers and non-rampers. During the delay period, ramper motor-units (*shown in olive*, *see Fig 2A)*, showed early changes in spiking activity (onset:∼-642ms; across population: -593.55±45.58ms, t(8)=-13.02, p<0.001) from the baseline. The slope or growth rate for the ramping activity (slope: 1.49 sp2/s; 3.32±0.99 sp/s^2^, t(8)=3.35, p=0.01) during the delay period was significant and it resulted in a higher level of activity at the time of GO cue (Δ2.80±.68sp/s, t(64)=4.062, p<0.001, across the population: Δ2.78±0.52 sp/s, t(8)=5.29, p<0.001) in the delayed task condition (as shown in bottom panel, Fig 2A). 41% of the units (9 out of 22; in 7 out of 9 subjects) showed a similar trend. In contrast, non-ramper motor units (*shown in dark green, see bottom panel Fig 2A*) showed no significant early recruitment or changes in activity during the delay time relative to the baseline (ΔActivity from baseline: Δ0.070±0.35sp/s, t(37)=0.20, p=0.841, across population: 0.23±.14 sp/s, t(12)=1.59, p=0.137). However, most of these units (13 out of 22, 59%) increased their activity following the GO cue and just prior to movement onset.

**Fig 2:**
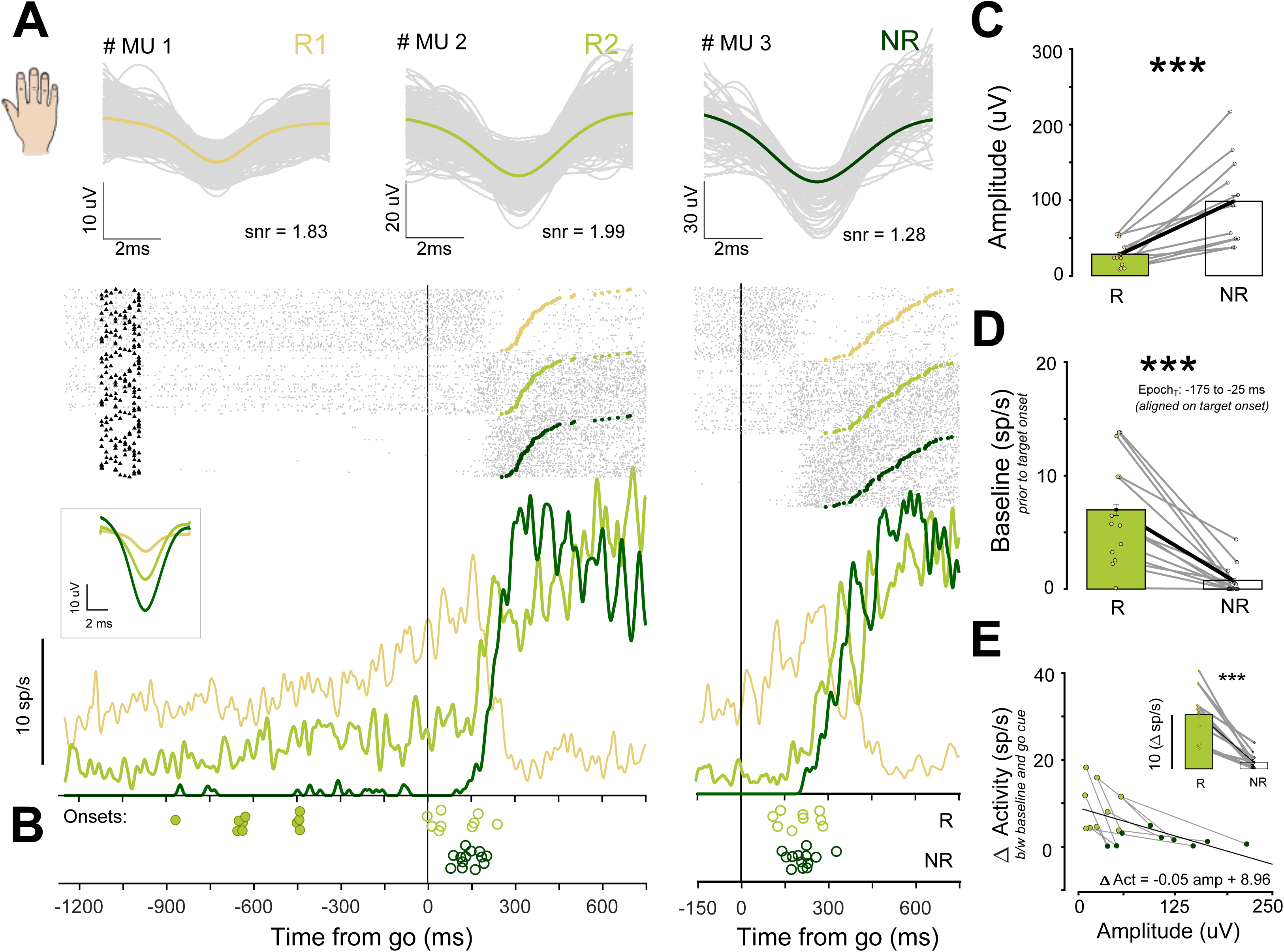
Recruitment of motor units based on amplitude size and baseline activity. A) *Ramper and non-ramper motor units:* Top Panel. Examples of motor spikes captured during a session. The activity profile for each motor unit was arranged based on spike amplitude size and is separated in the raster plot below. Each spike train in the raster plot represents a trial. Gray marks the time of occurrence for each spike. Filled triangles (in black) and circles (Skin and Olive: rampers, Green: non-rampers) show the time for target onset and initiation of movements following the GO cue, respectively. Bottom: Solid lines show the average activity for different motor units that were isolated for the representative session (Skin and Olive: rampers, Green: non-rampers). Inset shows the averaged spike waveforms across trials for the session. B) Filled and unfilled circles represent the onset time for rampers (olive) and non-rampers (green) before and after the GO cue, respectively. C) Bar plots showing the average amplitude of spike waveforms for rampers (skin) and non-rampers (green) from each session. D) Bar plots show the average basal firing rate prior to target onset for rampers (skin) and non-rampers (green) from each session. E) Recruitment pattern for small amplitude rampers (skin) and non-rampers (green). Each grey line represents the data collected from the same session. The black line represents the best fit linear regression model for the data set. The bar plots in the inset represent the average recruitment for rampers and non-rampers at the time of GO cue. * means P<0.05; ** means P<0.01; *** means P<0.001.

Since the SDF showed two phases of modulation; one prior to the GO cue (preparatory phase) and one after the GO cue (planning phase), we further characterised the differences in responses between rampers and non-rampers during the epoch between the GO cue and the movement onset (RT period) as well. The activity at the time of GO cue was lower for non-rampers when compared to rampers for both delayed (rampers: 9.18±1.82 sp/s; non-rampers: 1.99±0.48 sp/s, two sample t-test t-test: t(20) = 4.45, p<0.001) and immediate movements (rampers: 6.44±1.51 sp/s; non-rampers: 1.67±0.5 sp/s, two sample t-test: t(20) = 3.42, p=0.003). Interestingly, an earlier onset in activity was observed for rampers compared to non-rampers, following the GO cue during the delayed task condition (onsets: rampers: 94±28.10 ms; non-rampers: 139.53±10.9, two sample t-test: t(20) = 1.71, p=0.05). However, no such advantage, for rampers were observed during the immediate task condition (onsets: rampers: 196.3±22.07 ms; non-rampers: 208.15±13.19 ms, two sample t-test: t(20) = -0.48, p=0.63). Further, the growth rate or slope following the GO cue was significantly steeper for non-rampers when compared to rampers for both delayed (growth rate: rampers: - 17.18±.19.99 sp/s^2^; non-rampers: 76.01±12.24 sp/s^2^, two sample t-test: t(20) = 4.21, p<0.001) and immediate movements (growth rate: rampers: 2.79±18.24 sp/s^2^; non-rampers: 53.76±9.56 sp/s^2^, two sample t-test: t(20) = -2.69, p=0.014). In addition, the threshold level or the minimum activity reached by motor units prior to movement onset were significantly higher for non-rampers in comparison to rampers for both the delayed (threshold: rampers: 10.73±.2.48 sp/s; non-rampers: 19.97±3.09 sp/s, two sample t-test: t(20) = -2.16, p=0.042) and immediate (threshold: rampers: 11.18±2.55 sp/s; non-rampers: 19.65±2.93 sp/s, two sample t-test: t(20) = -2.05, p=0.05) task conditions.

We further tested whether motor-units follow recruitment patterns could be explained by Henneman’s principle, which posits that smaller motor units are recruited prior to larger motor units, under the assumption that, spike amplitude is a proxy of motor unit size. Consistent with Henneman’s principle we observed that rampers had significantly smaller spike waveform amplitudes (rampers: 28.45±5.54 µV, non-rampers: 98.39±18.06 µV, Pairwise t-test: t(10) = -4.60, p<0.001; **Fig 2C**) than non-rampers, which were recorded simultaneously. Second, rampers had a higher baseline firing rate) compared to non-rampers (rampers: 7.39±0.92 sp/s, non-rampers: 1.52±0.41 sp/s; Pairwise t-test: t(10) = 4.51, p<0.001; **Fig 2D**). Third, changes in activity during the delay time was negatively correlated (Linear regression: β=-0.05, p<0.001; Pearson’s r = -0.4971, p=0.02) to spike amplitude size (**Fig 2E**). Finally, the recruitment of motor units during the delay period based on spike amplitude size was negatively correlated to the increase in the activity from baseline at the time of GO cue. In other words, rampers showed more activity when compared to non-rampers (across population: rampers: 10.01±1.31 sp/s, non-rampers: 1.61±.46 sp/s; Pairwise t-test: t(10) = 5.65, p<0.001; result shown as inset in **Fig 2D**).

To summarise, a significant number of motor units (41%, 9 out of 22 units; in 7 out of 9 subjects and 33%, 11 out of 33 including supplementary data) showed modulations during the delay period. These motor units ‘rampers’ had smaller spike amplitudes, higher spontaneous activity and showed more modulation in activity prior to the GO cue when compared to the non-rampers (59%, 13 out of 22; 66%, 22 out of 33 including supplementary data) which showed modulations following the GO cue during the RT.

### Spatial and temporal information encoded during the early recruitment of motor unit activity

We first characterized the movement field (MF) of the muscle activity as the direction with maximum activity, -125 to 25 ms relative to movement onset (see, **<u>Fig S1</u>** and **<u>Fig S5</u>**). The average directional tuning across the population was approximately ∼104^0^ for the anterior deltoid muscle and indicated a right-upward component, as expected for a shoulder flexor (**<u>Fig S6</u>**). To calculate the time when motor activity could predict the direction of an upcoming movement, we divided the trials into 2 groups: trials where movement went into the MF (in-MF) and trials where the movements were made outside the MF (out-MF). An ROC analysis between in-MF and out-MF responses, for each time bin during visual, delay and movement epochs was performed by calculating the area under the curve (AUC) statistic which ranges from 0 (no separability) to 1 (complete separability). An AUC range from 0.5 to 1.0 indicates no separability to complete separability, respectively, for in-MF versus out-MF directions.

**Fig 3**, shows the spatio-temporal response profile for rampers. As observed for the motor unit activity shown in **Fig 3A**, different phases of modulations were observed during the delay epoch and reaction time epoch for the in-MF response compared to the out-MF direction for the delayed and immediate task (left vs right). AUC value showed an increasing trend (increase at GO cue: Δ 0.086 from baseline; onset: -235ms) during the delay period in the delay time task. **Fig 3B** shows this trend was consistent and could be seen across the population. The discriminability between in-MF and out-MF responses increased monotonically with time during the delay period between target onset to GO cue for rampers across the population (see inset, ΔAUC = 0.06±.008; t(8) = 7.39, p<0.001). The trend observed for the example unit was consistent across the population and onsets could be marked as early as -335.7±84.64ms prior to the GO cue (see methods). Even though the AUC values during the delay times were low (<0.7), discrimination times at these levels were observed for each subject.

**Fig 3:**
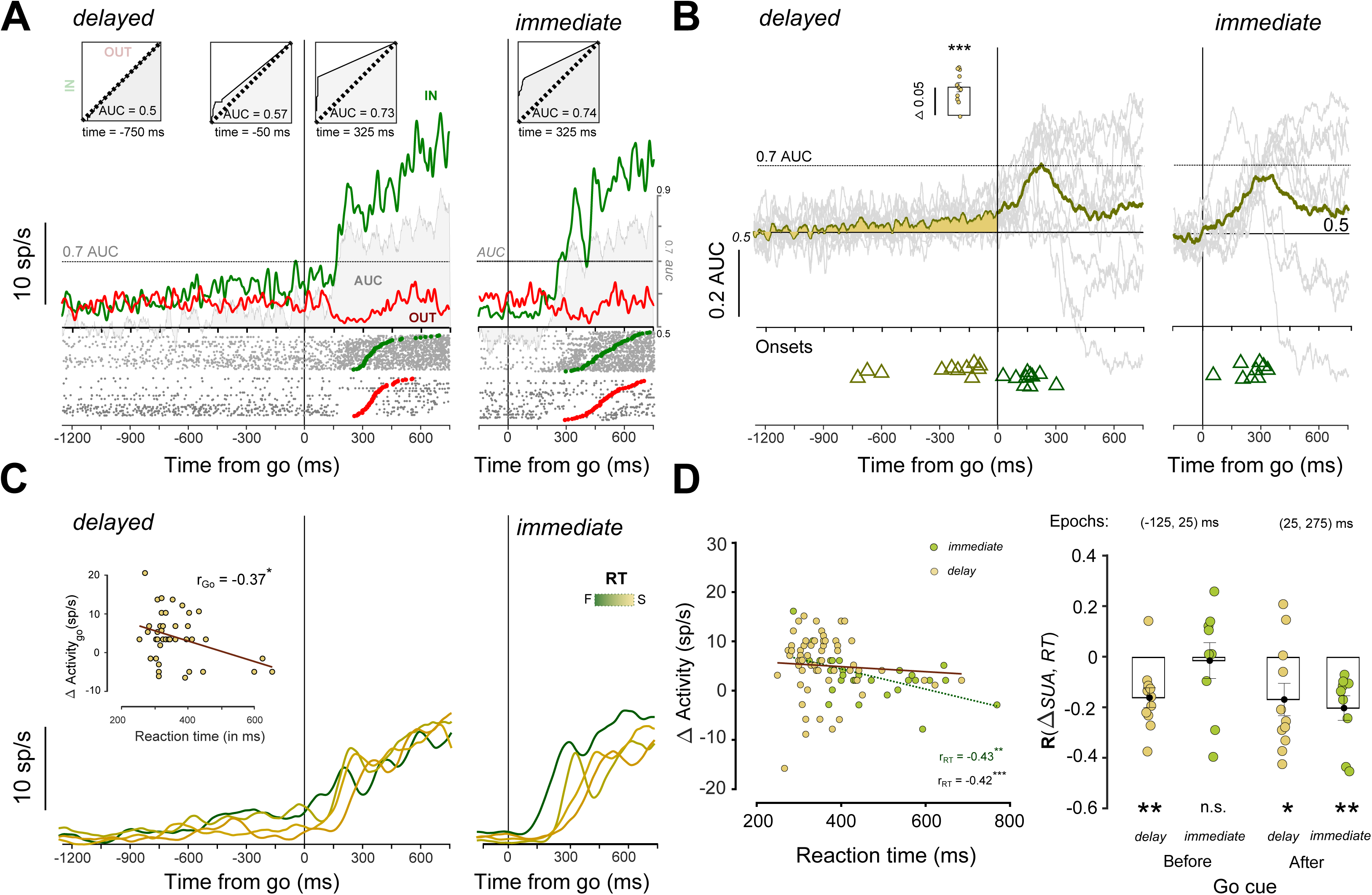
Rampers showing spatially selective response and temporal correlation for upcoming hand movements during the delay and reaction time periods. A) A representative session showing ramping motor unit activity for delayed and immediate hand movements, aligned on the GO cue for movements made towards (green) and away (red) from the movement field. The inset below shows the spiking activity of the motor unit for each trial marked in grey and the time when movements were initiated towards (green circles) or away (red circles) from the movement field. The grey shaded region represents the area under the curve (AUC) used for measuring the discriminability between in-MF and out-MF response for the EMG signal at each time bin. The inset above shows the separability between in-MF and out-MF response during baseline (−750ms), prior to the GO cue (−50ms), and following the GO cue (325ms) for the representative motor unit. B) Shown in grey are the AUC values from the ROC analysis for EMG responses in-MF and out-MF across different sessions for delayed (left) and immediate responses (right). The solid line (brown) shows the average mean value for the population. Inset. Bar plot for the average increase in AUC value from the baseline prior to the GO cue. Triangles (in olive) mark the onset for direction discriminability that gradually increases during the delay period for AUC values across all sessions. Triangles (in green) shows the direction discriminability onsets during the reaction time period based on an AUC>0.7 that demarcates a second later phase of recruitment. C) Activity for the ramping motor unit during delayed and immediate movements, aligned on the GO cue for different reaction times (RTs). Inset: Trial-by-trial scatter plot (olive) for change in activity from baseline prior to the GO cue as a function of RT. D) Left panel. Scatter plots of ramping activity following the GO cue for delayed (olive) and immediate movements (green) as a function of RT for the representative session. Right Panel. Bar plot for the Pearson’s correlations coefficient between change in activity from baseline prior to the GO cue and RT for delayed (olive) and immediate hand movements (green). *n.s.* means not significant; * means P<0.05; ** means P<0.01; *** means P<0.001.

Following the GO cue, discriminability was low, indicating that either the activity of these small amplitude size units could not be well isolated or were heterogenous in nature, in that some units interestingly, decreased their activity prior to movement onset. Nonetheless, a threshold value of 0.7 was used to mark the onset for the average direction discrimination time during the reaction time or planning phase. As expected, from the hold time effect, motor-unit activity showed differences in discriminability between in-MF and out-MF responses based on task conditions. For the delayed task, the AUC value reached the set threshold earlier when compared to immediate movements. (delayed task: 144±18.15 ms, immediate task: 242.88±28.3 ms; pairwise t test: t(8) = -4.61, p<0.001). A similar result due to the hold time effect was also observed for non-rampers (see *top panel*, **<u>Fig S8</u>**). In contrast to rampers, the activity of non-rampers for in-MF and out-MF conditions during the delay epoch, were similar and the AUC ∼0.5 across the population. The AUC value remained largely unchanged for the non-rampers until the GO cue appeared. Following the GO cue, the direction discriminability increased monotonically during the reaction time period. For the same threshold level (AUC=0.7),, an earlier onset in direction discrimination time was detected for the delayed task when compared to the immediate task (delayed task: 270.15±23.54ms; immediate task: 409.53±34.28ms; pairwise t-test: t(12) = -5.91, p<0.001), as was observed for the rampers., Taken together, we observed that spiking activity of the rampers provided early spatial information about upcoming movements when compared to non-rampers in both the delayed task (ramper: 144±18.15ms, non-ramper: 270.15±23.54ms; t-test: t(20) = -3.92, p<0.001) and more interestingly for immediate movements (rampers: 242.88±28.3ms, non-rampers: 409.53±34.28ms; t-test: t(20) = -3.499, p<0.001) even though their onset time for modulation were similar following GO cue.

Next, we looked at the correlations between the spiking activity and reaction time to study aspects of temporal information present within motor units during the delay period and reaction time epochs for delayed and immediate reaches. **Fig 3C** shows the response profile of motor activity for different RTs for a representative ramping unit. As rampers showed modulation during the delay period, a window of 125ms prior to GO cue and 25ms after the GO cue, was considered for the analysis to check if motor activity contained any temporal information about the upcoming movements. Consistent with this notion, we observed that the ramp up of activity during the delay time was negatively correlated with reaction time (r_Go_ = -0.367; p=0.01, see *inset* in **Fig 3C**). Furthermore, the negative correlations became stronger and significant at the single trial levels for both delayed and immediate hand movements between 175ms to 325ms following the GO cue for the representative motor-unit shown (delayed: r_RT_ = -0.42; p<0.001; immediate: r_RT_ = -0.43, p=0.018, see left panel **Fig 3D**). Right panel in *Fig 3D* summarises the results seen above across the population for ramping motor units which echoed the result observed in *Fig 3C*. For non-rampers as well, the negative correlations between activity and RT became stronger and significant for both delayed and immediate hand movements between 25ms to 275ms following the GO cue and was consistent across the population (delayed task: r_RT_ = -0.3±.06; t(12) = -4.81, p<0.001; immediate task: r_RT_ = -0.3±.07; t(12) = -4.08, p=0.0015, see bottom panel **<u>Fig S8</u>**).

To summarise, the activity in rampers during the delay period in the delay task could be used to discriminate the upcoming direction and RT of the movement. Additionally, the response of non-rampers, following the GO cue, could also discriminate the upcoming direction and RT of the movement.

### Single trial onsets and initiation of movements

In the previous section, we observed that the activity of rampers could be characterised by two phases of activity: an early preparatory phase prior to the GO cue and a later planning phase that occurred after the GO cue but prior to movement initiation. To test whether the two phases of activity could also be seen in single trials, a Poisson spike train analysis was used to mark the onset for changes in spiking activity for each trial (see methods for details). Since the distribution of the ISI’s were different for delay period and reaction time (see **<u>Fig S9</u>**), we analysed the trials in two separately epochs (i) prior to GO cue during the delay period and (ii) following GO cue during the reaction time period.

Shown for the example ramping motor unit in **Fig 4A** are the onsets for the spike train during the preparatory (in green triangles) and planning (in red) phase aligned on GO cue for trials sorted by reaction time. We observed that the onset times identified using the Poisson spike train analysis during the ramping or early preparatory phase were consistent with the AUC analysis and typically occurred between 400 and 50ms before the GO cue. These early onsets during the delay or preparatory period were significant and consistent across the population (onsets prior to GO cue: -335.46±25.03ms; t(8) = -12.99, p<0.001). Further, as shown in **Fig 4B** for the representative ramping motor unit, preparatory onsets for the spiking activity were negatively correlated to the baseline firing rate for the motor unit (baseline: r_bs_ = -0.31; p=0.03, see *left panel* **Fig 4B**) This result was significant and consistent across the population (baseline: r = -0.19±.07, t(8) = -3.39, p=0.01, see **Fig 4C**), suggesting that higher the baseline activity of the motor unit, the greater was the chance for detecting early changes in the spiking pattern for rampers. However, though such preparatory activity could be seen during the delay period, these changes at single trial level in rampers, were not predictive of the subsequent reaction time across the population (reaction time: r = -0.12±.08, t(8) = -1.43, p=0.2).

**Fig 4:**
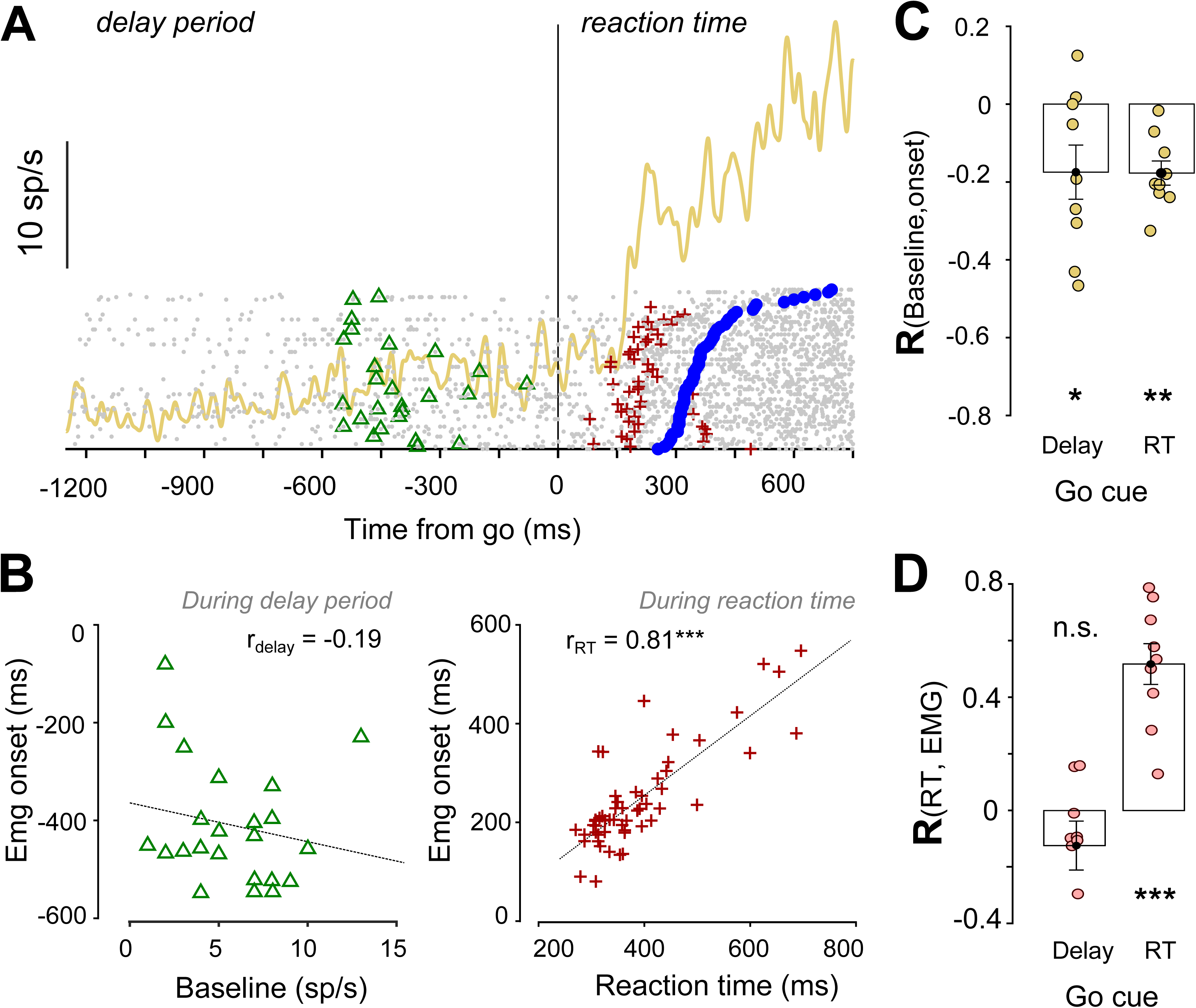
Single trials onsets for rampers during the delay and reaction time periods. A) Example showing different phases of ramping activity observed for the motor unit when aligned to the GO cue for the delayed reach movement. Gray dots in the raster mark the spike train for each trial sorted on RT. Green (triangles) and red (+, plus sign) markers represent the EMG onset, detected using Poisson spike train analysis for both delay and reaction time epochs, respectively. The blue (circles) markers show the onset of movements following the GO cue for each trial. B) Left Panel. Shows the scatter plot of EMG onsets detected during the delay period based on spiking activity for each trial as a function of baseline activity prior to the GO cue. Right panel. Scatter plot for EMG onsets detected during the reaction time epoch based on spiking activity for each trial as a function of reaction time. C) Bar plot shows the average Pearson’s correlation coefficient between the baseline activity of the motor unit and EMG onsets detected during the delay and reaction time periods for delayed movements. D) Bar plot shows the average Pearson’s correlation coefficient between reaction time and EMG onsets detected during the delay and reaction time periods for delayed movements. *n.s.* means not significant; * means P<0.05; ** means P<0.01; *** means P<0.001.

Next, we carried out the same analysis for the later planning phase that spanned the RT period as well. Onsets, were shorter for delayed movements compared to immediate movements (delayed: 270.33±15.59 ms, immediate: 368.74±12.88 ms) even at the single trial level, in accordance with the hold time effect, following the GO cue during reaction time. This result was consistent across the population as well (delayed: 171.93±20.45 ms; t(8) = 8.40, p<0.001, immediate: 244.28±33.08 ms; t(8) = 7.38, p<0.001). Like the delay period analysis, we tested whether there was any relationship between planning onsets and baseline activity and reaction time. As expected, even for the later planning phase, onsets were observed to be negatively correlated to baseline activity for the example ramping motor unit (delayed: r = -0.23, p=0.27 and immediate: r = -0.52, p=0.017) as well as across the population for both delayed and immediate movements (delayed: r = -0.17±.03, t(8) = -5.70, p<0.001 and immediate: r = -0.47±.04, t(8) = -11.88, p<0.001)., Onsets during the planning or RT period showed stronger positive correlations with the time it took to initiate movement following GO cue in both the example ramping motor unit (delayed: r = 0.67, p<0.001 and immediate: r =0.78, p<0.001; see *right panel* **Fig 4B**) and across the population as well (delayed: r = 0.51±.07, t(8) = 7.18, p<0.001, immediate: r = 0.55±.06, t(8) = 9.60, p<0.001, see **Fig 4D**).

We also carried out the analysis for the RT period for non-rampers as they show activity post GO cue (see *top panel*, **<u>Fig S12</u>**). Onsets during the reaction time showed a similar trend as the late ramping activity; with shorter onset times for delayed movements (onsets: 308.47±14.03 ms; t(12) = 21.97, p<0.001) when compared to immediate movements (onsets: 393.69±11.56 ms; t(12) = 34.03, p<0.001). Similar to the rampers, onsets during the planning phase were negatively correlated to baseline activity (delayed: r = -0.22±.09, t(12) = -2.50, p=0.02, immediate: r = -0.32±.08, t(12) = -2.47, p=0.029) and positively correlated to the reaction time (delayed: r = 0.57±.08, t(12) = 6.73, p<0.001, immediate: r = 0.57±.10, t(12) = 5.28, p<0.001) for both delayed and immediate movements across the population.

To summarise, we carried out the single trial analysis for ramper and non-ramper motor units using Poisson spike train analysis Interestingly, the early preparatory phase onsets were correlated to baseline activity but not with reaction time while the onsets of the second planning phase were more strongly correlated to reaction time.

### Motor unit activity can be described by accumulate to threshold models of response initiation

We tested whether an accumulation to threshold model could describe the initiation of movements by analysing the response of rampers and non-rampers. In previous studies (Hanes and Schall 1996) such ramping activity during reaction time has been formalized in accumulator models which has been used only in the context of immediate movements to understand the neural basis of response preparation. Here, we tested if these parameters describing accumulation could be extended to understand initiation of reach movements in the delay period as well. Responses divided into *fast and slow reaction times* (see methods). Different parameters of the accumulator model-(i) activity at the time of GO cue (ii) slopes and (iii) onset for activity during the delay period (iv) and threshold i.e., motor activity at the time of movement initiation were calculated motor units under both the conditions (see method). **Fig 5A** shows the response of a ramper from a representative session for fast and slow RTs during delayed movements (*Fig5A*, top left panel) and movement onset (F*ig5A*, bottom left panel) and the same for immediate movements (*Fig5A*, right top and bottom panels).

**Fig 5:**
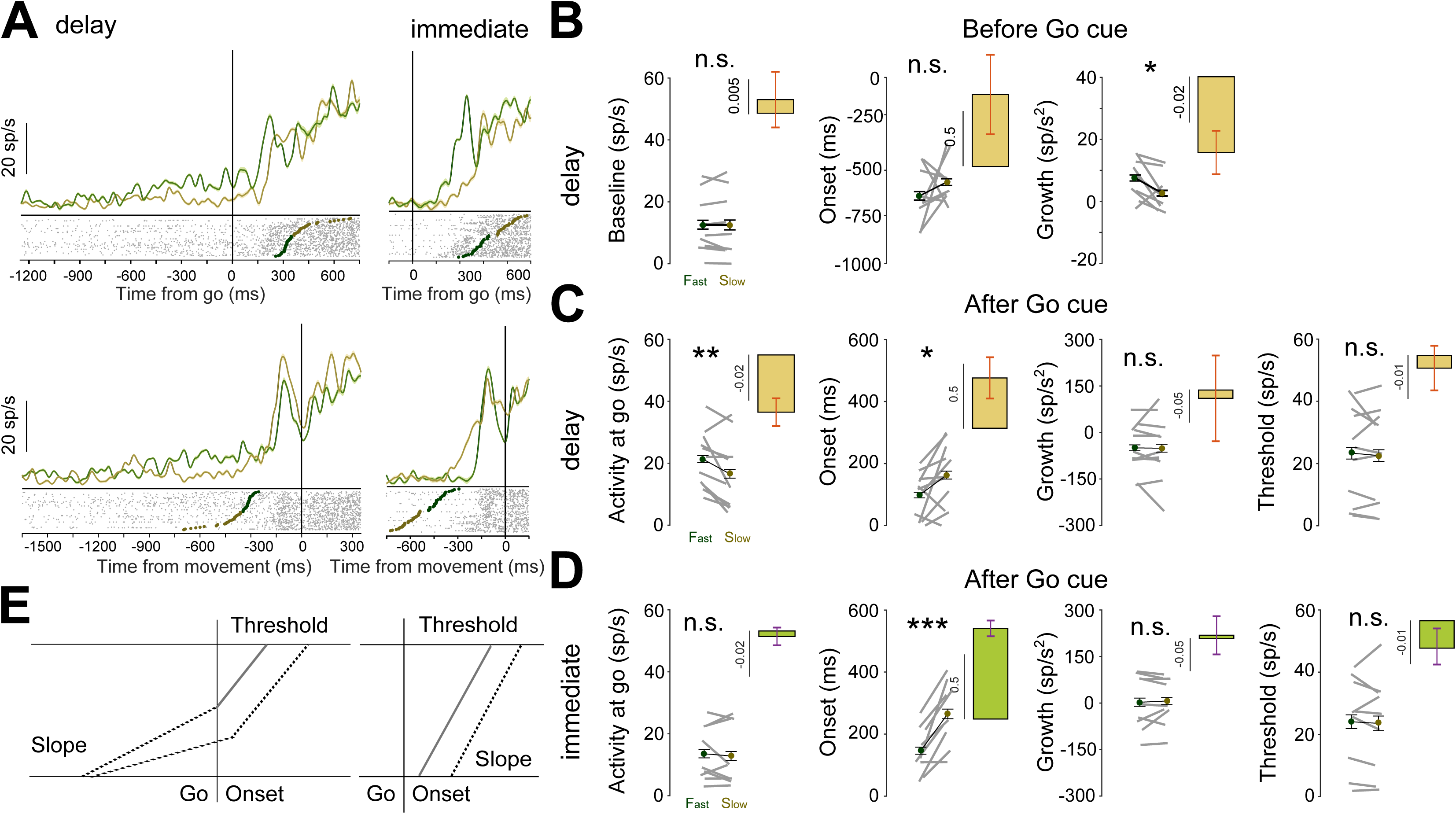
Ramping activity can be described as an accumulator model. A) Activity for a ramping motor unit aligned on the GO cue (top panel) and movement onset (bottom panel) in the delayed and immediate reach tasks. B) The bar plots show the change in motor activity during the baseline (first panel), for onset (second panel), and growth rate (third panel) prior to the GO cue for slow (olive) and fast (green) reaction times. C) Same parameters from B that were used to capture changes in motor activity were used to measure changes at the GO cue (first panel), onset (second panel) and growth rate (third panel), and threshold (fourth panel) following the GO cue. Each line in grey represents the motor unit that was obtained across subjects. D) same as the lower panel of delayed but for the immediate reach task. E) Schematic representation of the accumulator model that summarizes the data for rampers during delayed and immediate movements. *n.s.* means not significant; * means P<0.05; ** means P<0.01; *** means P<0.001.

We observed that the activity of rampers at baseline during fast and slow RTs under the delayed condition were similar (delayed: fast RT: 12.60±3.20 sp/s, slow RT: 12.52±.3.52 sp/s; t(8) = 0.12, p=0.90; first panel in **Fig 5B**). There was also no significant change in onset times during the delay period for spiking activity observed for fast and slow RTs (delayed: fast RT -549.6±75.59 ms; slow RT -421.2±82.8 ms; t(8) = 1.73, p=0.06; second panel in **Fig 5B**). This observation recapitulates the lack of any relation between early onsets and reaction time that was described in the single trial analysis (Fig 4). However, as shown in fig 3, we observed that during the delay period, there was significant increase in growth rate for rampers for fast RTs compared to slow RTs (delayed: fast RT: 7.55±2.18 sp/s, slow RT: 2.66±1.97 sp/s; t(8) = 2.37, p=0.02; third panel in **Fig 5B**).

Next, we analysed changes in the activity of rampers following the GO cue for fast and slow RTs under both delayed and immediate task conditions. For the delayed but not the immediate task, the activity around the GO cue was higher for faster reaction times compared to slow RTs. This observation was consistent across the population (delayed: fast RT: 20.43±3.29 sp/s, slow RT: 16.43±.4.07 sp/s; t(8) = 2.83, p=0.02; first panel in **Fig 5C**; immediate: fast RT: 13.54±2.96 sp/s, slow RT: 12.85±3.23 sp/s; t(8) = 0.55, p=0.59; first panel in **Fig 5D**). We found a significant increase in onset times for ramping activity during fast and slow RTs for delayed as well as immediate movements (delayed: fast RT 126.1±15.76 ms; slow RT 158.7±23.8 ms; (t(8) = -1.79, p=0.06; second panel in **Fig 5C**; immediate: fast RT: 156.3±22.14 ms and slow RT 274.7±30.13 ms; (t(8) = -6.38, p<0.001; second panel in **Fig 5D**). No significant difference in the growth rate for the spiking activity from the time of GO cue was observed for movements with fast and slow RTs across the population for both delayed as well as immediate movements (delayed: fast RT: 20.12±98.77 sp/s, slow RT: 0.54±55.7 sp/s; t(8) = 0.37, p=0.36; third panel in **Fig 5C**; immediate: fast RT: -32.9±64.5 sp/s, slow RT: -44.2±57.8 sp/s, t(8) = 0.59, p=0.28; third panel in **Fig 5D**). In addition, we also aligned the response on movement onset and tested whether ramping activity reached a fixed level or threshold prior to initiating movements for both fast and slow RTs during delayed and immediate task conditions. We observed that the threshold activity in both the tasks, under delayed and immediate conditions, was similar for slow and fast RTs. This observation was consistent across the population (delayed: fast RT: 26.4±3.61 sp/s, slow RT: 27.02±4.74 sp/s; t(8) = -0.32, p=0.75; fourth panel in **Fig 5D**; and immediate: fast RT: 26.32±3.9 sp/s, slow RT: 27.54±.4.6 sp/s; t(8) = -1.09, p=0.30; fourth panel in **Fig 5D**).

Next, we tested whether the same parameters could be used to explain the motor activity pattern for non-rampers following the GO cue for delayed and immediate movements. **Fig 6A** shows the response from a representative non-ramper from the same session for fast and slow RTs during delayed and immediate movements aligned on the GO cue (top panel) and movement onset (bottom panel). For the delayed and immediate conditions no change in baseline motor activity was observed around the GO cue for fast and slow RTs and this observation was consistent across the population (delayed: fast RT: 4.72±1.2 sp/s, slow RT: 3.48±0.9 sp/s; t(12) = 1.90, p=0.08; first panel in **Fig 6B**; immediate: fast RT: 3.17±0.9 sp/s, slow RT: 3.28±1.03 sp/s; t(12) = -0.32, p=0.75; first panel in **Fig 6C**). Interestingly, we found significant increase in onset times for fast and slow RTs for the delayed and immediate task conditions as well (delayed: fast RT: 103.84±6.43 ms and slow RT: 160.77±16.35 ms, t-test: t(12) = -4.11, p<0.001, second panel in **Fig 6C**; immediate task condition: fast RT: 163.46±13.97 ms and slow RT: 212.38±11.2 ms, t(12) = -3.6, p=0.001; second panel in **Fig 6D**). Also, we observed significant and consistent decrease in growth rate across the population for fast and slow RT for movements made under delayed and immediate conditions (delayed: fast RT: 203.85±39.9 sp/s and slow RT: 153.06±23.48 sp/s; t-test: t(12) = 2.04, p=0.032, third panel in **Fig 6C**; Immediate: fast RT: 165.41±29 sp/s and slow RT: 212.38±11.2 sp/s; t-test: t(12) = 5.86, p<0.001; third panel in **Fig 6D**). Like the rampers, we also aligned the population response on movement onset and tested whether threshold activity at the time of movement onset remained same for fast and slow RTs during delayed and immediate task condition. Activity in both the tasks under delayed and immediate conditions was similar for slow and fast RTs. This observation was consistent across the population (delayed: fast RT: 30.38±6.3 sp/s, slow RT: 30.02±6.2 sp/s; t(12) = 0.30, p=0.76; fourth panel in **Fig 6C**; immediate: fast RT: 29.82±6.25 sp/s, slow RT: 30.66±6.37 sp/s; t(12) = -0.77, p=0.45; fourth panel in **Fig D**).

**Fig 6:**
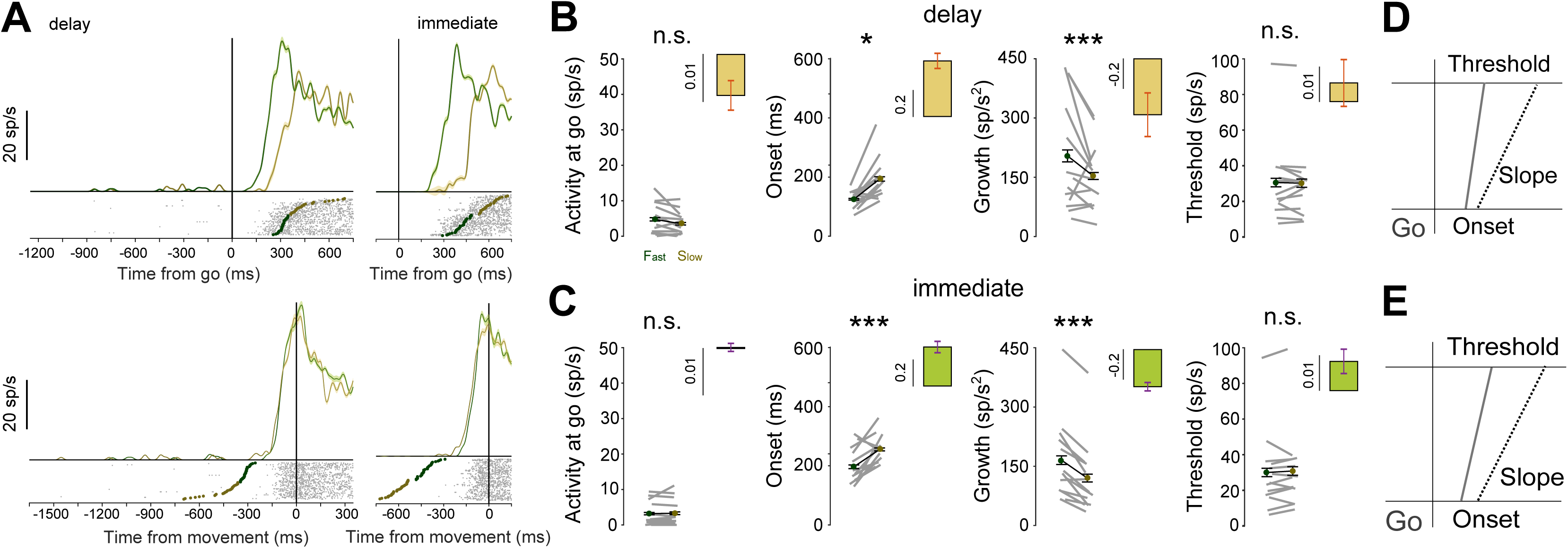
Non-ramping activity can be described as an accumulator model. A) Activity for a non-ramping motor unit aligned on the GO cue (top panel) and movement onset (bottom panel) for the delayed and immediate reach conditions. B) The bar plots show the average change in motor activity firing rate at the GO cue (first panel), onset (second panel), growth (third panel), and threshold (fourth panel) for slow (olive) and fast (green) reaction times. Each line in grey represents the data that was obtained across each session. C) same as B but for the immediate reach task. D) Schematic representation of the accumulator model following the GO cue that summarizes the data for non-rampers during the delayed condition E) Same as D but for immediate movements. *n.s.* means not significant; * means P<0.05; ** means P<0.01; *** means P<0.001.

In summary, by analysing the data for movements made with delayed and immediate movements, we tested the applicability of the linear accumulator model and found that different parameters/architectures explained the modulation of motor unit activity during the reaction time for both rampers (**Fig 5E**) and non-rampers **(Fig 6E and F)**. Specifically, activity at the GO cue, growth rate and onsets played an important role during the reaction time period to help initiate an upcoming movement.

### Experiment 2: Isometric force driven cursor movements

To further test our results, particularly in relation to previous studies that have not shown early preparatory activity during the delay period, we also recorded from 16 subjects during an isometric force task which involved cursor movements (see methods, for details), as was previously done by (Prut and Fetz 1999). In this task, 11 sessions were performed where delayed and immediate tasks were interleaved while an additional 5 subjects performed the delayed task in separate sessions. Similar to the hand experiment, delayed isometric movements had shorter reaction times compared to immediate isometric movements (see **Fig 1B**; delayed task: 377.83±9.99ms and immediate: 492.63±24.88ms; t(10) = -6.17, p<0.001). These results indicate that the delay period conferred a reaction time advantage. Also, like the hand experiments, high-density surface EMG signals were analysed using both conventional and raster-based method. Both the approaches gave comparable results (**Fig 1D**; Cursor: delayed task: Pearson’s r = 0.83 ± 0.05, p<0.0001 and immediate task: Pearson’s r = 0.70 ± 0.04, p<0.0001) and since the latter provided information about the spike waveforms in addition to the frequency of spike occurrence, results from raster-based approach are shown below, unless otherwise specified.

We identified 20 motor units (29 including additional 5 subjects’ data) in the data set by analysing one of the differential channels in the array per subject for this study. In contrast to the hand task, none of the motor units showed any significant modulations during the delay period (activity at GO cue: -Δ0.64±0.58sp/s, t(40)=-1.11, p=0.272; across population: - Δ0.08±0.28sp/s, t(19)=-0.30, p=0.77). Only one subject out of eleven showed a positive difference in activity during the delay time from baseline. **Fig 7A** shows the response of motor units captured during a typical session. Modulations in these units were observed following the GO cue during the reaction time epoch only. An earlier onset of motor activity was seen following the GO cue during the reaction time period for the delayed task condition when compared to the immediate task condition (onsets: delayed task: 134ms for the example session; across population: 124.5±17.28ms and immediate task: 343ms for the example session; across population: 186.1±25.46ms, pairwise t-test: t(19) = -1.90, p=0.036).

**Fig 7:**
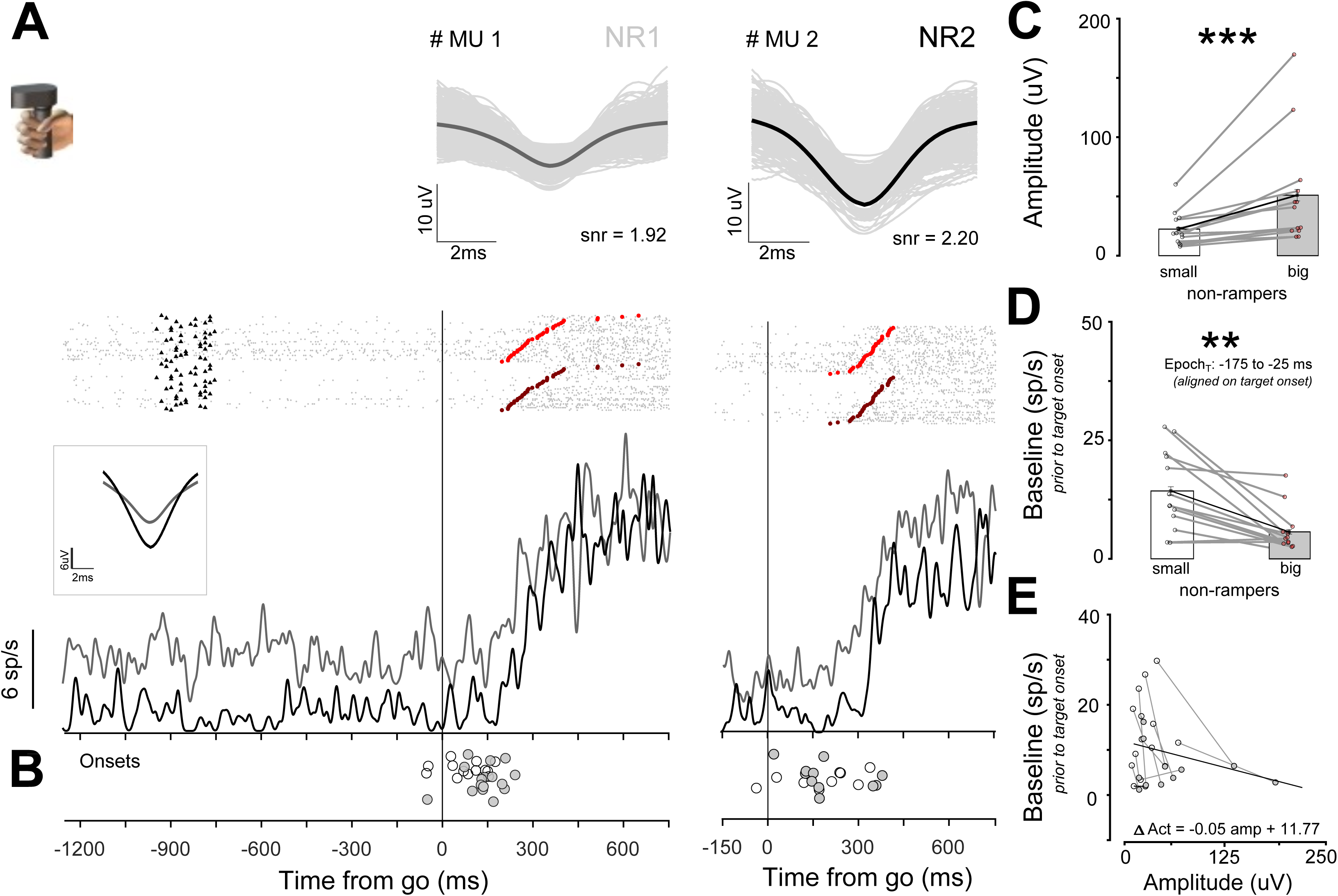
Recruitment of motor units based on amplitude size and baseline activity in the isometric cursor task. A) Top Panel. Example motor units captured during a session. The activity profile for each motor unit was arranged based on average spike amplitude size and is shown below in the raster plots. Each spike train in the raster represents a trial. Gray marks the time of occurrence for each motor unit spike from the time of the GO cue. Filled triangles and circles (red: small, brown; large; amplitude non-rampers) show the time for target onset and initiation of movements following the GO cue, respectively. Bottom: Solid lines show the average activity for different non-ramping motor units that were isolated for the representative session (grey: small, black: large). The inset shows the averaged spike waveforms pooled across trials for the session. B) Filled and unfilled circles represent the onset time for small (grey) and large amplitude (black) non-rampers before and after the GO cue, respectively. C) Bar plots showing the average amplitude of spike waveforms for small (grey) and large (black) non-rampers from each session. D) Bar plots show the average basal firing rate for small (grey) and large (black) amplitude size non-rampers from each session. E) Baseline activity for small (grey) and large (black) amplitude non-rampers. Each grey line represents the data collected from the same session. The black line represents the best-fit linear regression model for the data set. The bar plots in the inset represent the average recruitment for the rampers and non-rampers at the time of GO cue. * means P<0.05; ** means P<0.01; *** means P<0.001.

We further tested whether these motor-units followed a recruitment pattern similar to the hand experiments based on Henneman’s principle. First, for sessions with multiple non-rampers, we observed small and large spike amplitude motor units (non-rampers: small: 22.44±4.0 µV, large: 51.07±12.75 µV, Pairwise t-test: t(12) = -3.09,p=0.004; **Fig 7C**). Second, small amplitude motor-units had a higher baseline firing rate (non-rampers: small: 14.34±2.33 sp/s, large: 5.71±1.27 sp/s; Pairwise t-test: t(12) = 4.16, p=<0.001; **Fig 7D**) compared to large amplitude motor units. Third, baseline activity during the delay time was negatively correlated (Linear regression: β=-0.05, p<0.001; Pearson’s r = -0.4971, p=0.02) with the amplitude size of the spike waveforms captured during the same session. **Fig 7E**.

We next characterised the activity for non-rampers based on their responses during the reaction time period for the delayed and immediate task conditions. We observed that motor activity when aligned on movement onset reached a particular threshold prior to initiation of movements. This threshold value was similar for both the delayed and immediate task condition (Threshold for cursor movements: delayed task: 18.42±2.53 sp/s, immediate task: 17.38±2.52 sp/s; paired t test: t(19) = 1.52, p=0.072). Also, the growth rate of motor activity for delayed when compared to immediate task conditions was similar following the GO cue for cursor movements (growth rate: delayed task: 28.97±3.97 sp/s^2^, immediate task: 18.64±3.19 sp/s^2^; paired t test: t(19) = 2.69, p=0.007).

Activity during the delayed epoch was comparable and showed no trend or any modulation (at GO cue: Δ -0.01) from the baseline (**Fig. S11**). The AUC or separability between the in-MF and out-MF responses for non-ramping motor units was almost 0.5 and not significant. *Fig S11B, right panel* summarizes the results across the population for the non-rampers. The AUC value remained largely unchanged for the delay period until the GO cue appeared (non-ramper: ΔAUC = 0.004±.01; t(19) = -0.35, p=0.73). Following the GO cue, the direction discriminability increased monotonically during the reaction time. For the same threshold level (AUC=0.7), earlier onsets for direction discrimination time was observed for the delayed task condition when compared to the immediate task condition (delayed task: 333.85±21.27ms; immediate task: 466.9±33.95ms; pairwise t-test: t(19) = - 4.41, p<0.001). These results suggest spatial information reached periphery during the reach time epoch just prior to making an upcoming movement, consistent with prior work, but in contrast to the delayed reach task.

To test whether motor activity could explain the variability in reaction time (RT), we calculated a Pearson’s correlation coefficient on trial-by-trial basis between motor unit activity and reaction time, both before and after the GO cue, for delayed and immediate cursor movements. No correlation was seen during the delay time for cursor movements (**<u>Fig</u> <u>S12</u>**). The negative correlations became stronger and significant at the single trial level for both delayed and immediate cursor movements between 75 ms to 375 ms following the GO cue. *Fig S12D, left panel* shows a correlation plot for the representative session during reaction time (delayed: r_RT_ = -0.42; p=0.021; immediate: r_RT_ = -0.207, p=0.288). *Fig S12D*, *right panel* summarises the results across the population for the cursor experiment. As non-rampers showed no modulation during the delay time, the activity prior to the GO cue was not correlated with reaction times (delayed: t(19) = -0.015, p=0.98; immediate: t(19) = -0.75, p = 0.45). Also, as expected, prior to movement onset, the correlations with reaction time grew stronger for both delayed and immediate movements (delayed task: r = -0.2±.05; t(19) = -3.94, p<0.001; immediate task: r = -0.17±.04; t(19) = -3.82, p=0.0012). To summarise, for cursor movements since no significant modulation was seen during the delay period, behaviourally relevant spatial and temporal information was not present. However, and not surprisingly, the build-up in muscle activity could be used to discriminate the upcoming direction of movement during reaction time period.

Next, we carried out the single trial spike train analysis for the reaction time epoch for non-ramping motor units (see bottom panel, **<u>Fig S12</u>**). Interestingly and contradictory to the hand movement task, we observed that EMG onsets following the GO cue were not significantly different for both delayed as well as immediate conditions (delayed: 257.13±11.75 ms; immediate: 279.76±19.73 ms; t(38) = -0.98, p=0.33). Therefore, EMG onsets alone could not explain the delay time effect or difference in reaction time between the two conditions. Also, they were negatively correlated to baseline activity across the population (delayed: r = -0.19±.07, t(19) = -2.43, p=0.0251, immediate: r = -0.33±.05, t(19) = -6.48, p<0.001) for both delayed and immediate movements. Whereas EMG onsets during the reaction time were positively correlated with the time it took to initiate cursor movements. This result was consistent across the population for both delayed and immediate movements (delayed: r = 0.37±.10, t(19) = 3.58, p=.002, immediate: r = 0.26±.05, t(19) = 5.14, p<0.001), suggesting that an increase in firing rate plays an important role in initiating cursor movements.

We also tested whether the parameters of the accumulator model could be used to explain the motor activity pattern for initiation of cursor movements during delayed and immediate conditions. Interestingly, both onsets and growth rate were important factors and played an important role in initiating cursor movements. **Fig 8A** shows the response from a representative cursor session for a delayed and immediate task, for slow and fast RTs aligned on GO cue (top panel) and movement onset (bottom panel). For both the delayed and immediate conditions the motor activity around the GO cue for slow and fast RTs was not significantly different and this observation was consistent across the population (delayed: fast RT: 21.47±4.36 sp/s, slow reaction time: 22.59±4.43 sp/s; t(28) = -0.72, p=0.47; first panel in **Fig 8B**; immediate: fast RT: 25.41±4.71 sp/s, slow RT: 25.96±4.50 sp/s; t(19) = -0.49, p=0.62; first panel in **Fig 8C**). We observed a significant increase in onset times for fast and slow RTs for delayed and immediate conditions (delayed: fast RT: 89.38±23.94 ms and slow RT: 170.79±31.98 ms, t-test: t(28) = -2.61, p=0.007, second panel in **Fig 8B**; immediate: fast RT: 160.35±20.25 ms and slow RT: 244.20±31.97 ms, t(19) = -2.45, p=0.01; second panel in **Fig 8C**). Also, a significant decrease in growth rates for fast and slow RTs were observed during delayed and immediate conditions (Delayed: fast RT: 55.35±6.19 sp/s and slow RT: 38.41±6.76 sp/s; t-test: t(28) = 2.21, p=0.017, third panel in **Fig 8B**; Immediate: fast RT: 46.62±5.97 sp/s and slow RT: 29.10±5.77 sp/s; t-test: t(19) = 3.75, p<0.001; third panel in **Fig 8C**). In addition, we also aligned the response of motor units on movement onset and tested whether threshold activity at the time of movement onset differed for fast and slow RT during the delayed and immediate task conditions. Activity around movement onset for both the delayed and immediate conditions was not significantly different for slow and fast RTs. This observation was consistent across the population (delayed: fast RT: 25.41±4.77 sp/s, slow RT: 28.07±4.79 sp/s; t(28) = -1.54, p=0.13; fourth panel in **Fig 8B**; immediate: fast RT: 28.46±4.74 sp/s, slow RT: 30.07±4.83 sp/s; t(19) = -1.22, p=0.24; fourth panel in **Fig 8C**). To summarise, we observed that different parameters/architectures could explained the modulation of motor activity during the reaction time for both delay and immediate isometric movements (see, **Fig 8D**). Both, growth rate and onsets played an important role during the reaction time to help initiate isometric force driven cursor movements.

**Fig 8:**
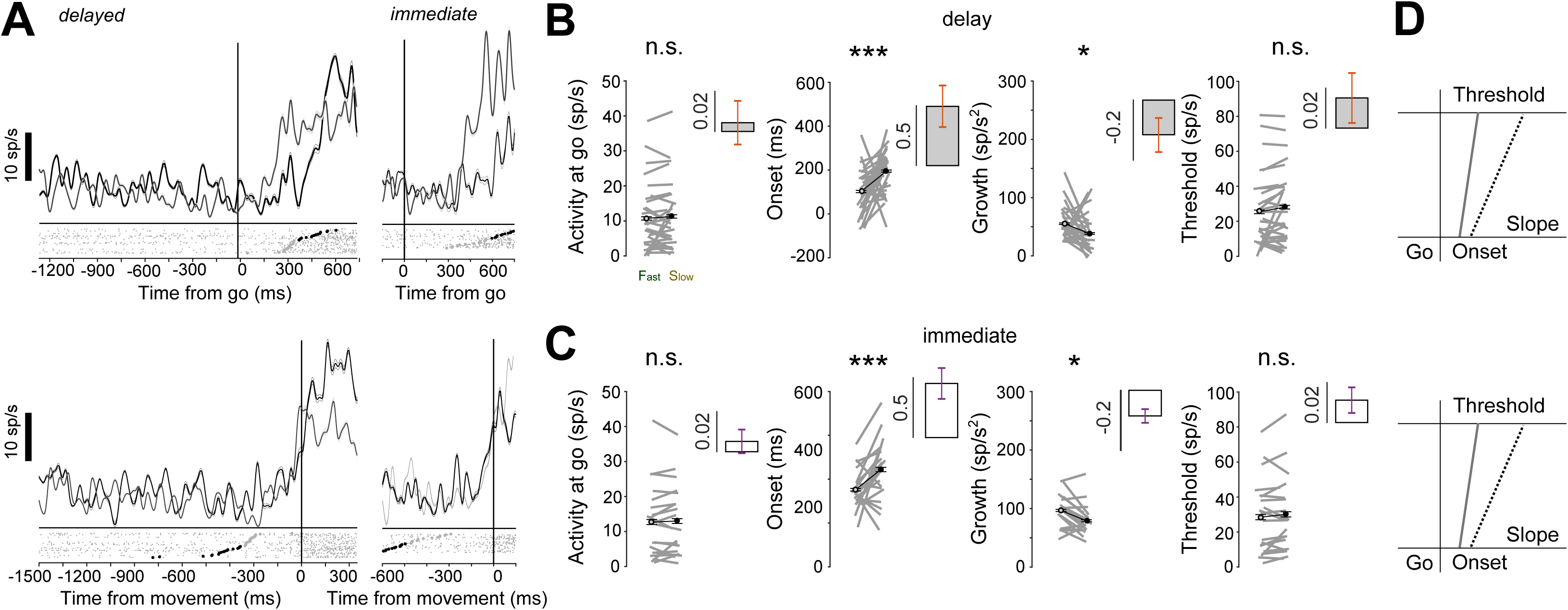
Isometric force driven initiation of cursor movements can be described by an accumulate to threshold model. A) Activity for a non-ramping motor unit aligned on the GO cue (top panel) and movement onset (bottom panel) for the delayed and immediate conditions. B) The bar plots show the average change in motor activity at the GO cue (first panel), onset (second panel), growth (third panel), and threshold (fourth panel) for slow (black) and fast (grey) RTs. Each line in grey represents the data that was obtained across each session. C) same as B but for the immediate reach task. D) Schematic representation of the accumulator model following the GO cue that summarizes the data for non-rampers during delayed (top) and immediate (bottom) movements. *n.s.* means not significant; * means P<0.05; ** means P<0.01; *** means P<0.001.

## Discussion

Our study is the first attempt at using high-density surface EMG to show the presence of early spatially specific recruitment of muscle activity that is also task dependent. Our results show both spatial and temporal information is encoded in motor activity during the delay period for hand movements but not for isometric cursor movements. However, after the GO cue, irrespective of the type of movement (hand or cursor), the activity during the reaction time period could be used to predict the variability in movement onset with respect to delayed and immediate tasks. Taken together, our results may explain the divergent observations regarding the presence or absence of early motor unit activity during the planning of reach movements in humans. Finally, we showed that an accumulation to threshold model, that has provided a simple framework to understand the initiation of saccadic movements, can also be used as a linking hypothesis to test the relation between the motor unit activity and the initiation of reach movements and their dependence on task context.

### Context dependent early recruitment of motor activity

A major unresolved issue pertaining is whether motor preparation is reflected in muscle activity during an instructed delay task, as has been observed for motor cortical neurons. Seemingly different results for the same task performed but under slightly different conditions have been observed by different groups. Prut and Fetz (1999) recorded from distal muscle groups while macaques were involved in an isometric force task, and they showed no modulation of EMG activity in muscles during the delay period despite activity being present in spinal cord neurons. However, while macaques were involved in making actual flexion-extension arm movements, Mellah et al. 1990 observed motor units were recruited during the delay period in proximal muscle groups. More recently, in humans, Papaioannou and Dimitriou 2021, also observed preparatory signals in Ia afferent but in the absence of changes in EMG activity. These disparate results could be a consequence of the different muscles used since proximal versus distal muscle groups have different delays in muscle activation which contribute partly to the observed transport delay between reach and grasp actions observed in overt behaviour (Jeannerod 1981). Thus, differences in proximal versus distal muscles may be a consequence of these muscle groups being subject to different levels of spinal inhibitory control.

In addition to different muscle groups, our results show an unexpected heterogeneity in the activity of EMG responses as well, in that delayed period activity is present in only small amplitude size motor units. Although in principle the spike amplitude can be conflated with the distance of a motor unit from the electrode, such early recruitment of small amplitude motor units is consistent with Henneman’s recruitment principle based on the size of the motor units. That being the case, our results suggest that such early recruitment may reflect the generation of small forces that cannot overcome the inertia to cause overt movements of the hand but still could facilitate reaction times in conjunction with the later recruitment of larger motor units.

However, the absence of such early recruitment in the isometric task is hard to explain since we observed no preparatory activity despite the deltoid muscle being subject to forces that were comparable to those generated for reaching movements (see **<u>Fig S2</u>**). Although this result requires further systematic study, we cautiously interpret these results as suggesting that the absence of early recruitment in isometric force-driven cursor movements might be a consequence of the absence of explicit kinematic planning. These results also align with Papaioannou and Dimitriou 2021; Prut and Fetz 1999, since their task was also cursor based and may not have necessitated explicit kinematic planning. In contrast, dynamics related activity of motor units that dominate EMG activity, is likely to be subjected to stronger inhibitory control. However, we cannot rule out alternative interpretations. Since we did not measure the positions of the joints, we cannot rule out their movements in the null space which would not affect end point position as a consequence of redundancy. However, this view fails to explain the absence of preparatory activity in the cursor task, which should have produced similar preparatory joint activity.

### Early recruitment of muscle activity encodes spatial and temporal information

Previous studies in psychology have suggested that information can flow in a continuous mode from successive processing stages (Mcclelland 1979; Meyer et al. 1988). Similarly, studies in the oculomotor and motor systems have suggested that information can flow in a continuous mode from different brain areas that are involved in visual to movement transformations (Bichot et al. 2001; Cisek and Kalaska 2005; Crammond and Kalaska 2000; Schmolesky et al. 1998). We extended these ideas to study the flow of information into the peripheral musculature. Previous efforts have tried to address this issue in the context of immediate movements and have shown evidence of a stimulus linked response (SLR) in proximal arm and neck muscles (Corneil and Munoz 2014; Pruszynski et al. 2010), that may be associated with neural activity associated with the reticulospinal tract (Kearsley et al. 2021). However, since the SLR is mostly associated with reflexive movements, dissociating early and late components of movement planning from execution are challenging. By separating processes pertaining to *where* and *when* an action is to be made by using a delay period in the visually guided delay time paradigm, we now show that slower modulations in the EMG signal carried both spatial and temporal information about a prospective movement much before the GO cue, independent of the occurrence of an SLR.

The presence of both early and late components of motor planning could be seen in the AUC and temporal correlations (**Fig 3 and 4**). Whereas during the delay period, the AUC and temporal correlations with reaction times were modest, they dramatically increased after the GO period. This is not surprising, given the presence of variability after the GO cue is expected to dilute the contributions of preparatory activity to reaction times. This effect was also seen at the level of single trials where we observed robust correlations between the onset of EMG activity and reaction times in the planning period but not in the preparatory period. Nevertheless, what was surprising is the absence of such early modulations for cursor movements only. Such specificity with respect to task context suggests that motor unit recruitment is much more flexible than previously appreciated (Marshall et al. 2022; Mottram et al. 2005).

Although the specific role of such early muscle activity is unclear, such early muscle activity may allow the shoulder muscles to overcome the larger inertia of the limb to facilitate the initiation of hand movements. Such early activity could therefore either prime the proximal muscle groups, or drive or even control the subsequent dynamics related muscle activation as envisioned in equilibrium control models (Dimitriou and Edin 2010; Latash 2018; Polit and Bizzi 1979). Further, if such early activity is related to kinematics, it is expected to be spatially specific, as we observed for the agonist muscle, but not when the same muscle is an antagonist. The rather early activity observed in the absence of overt movement further suggests that this activity is likely to reflect the activation of small motor units, which may explain why their contribution in muscle EMG activity may have gone unnoticed.

### Assessing initiation of movements with an accumulator model that tracks motor activity recruitment

A Linear Accumulation to Threshold with Ergodic Rate (LATER) model has been used to successfully account for reaction time data in eye and hand movement tasks (Asrress and Carpenter 2001; Carpenter and Williams 1995; Dean et al. 2011; Gopal and Murthy 2015; Jana et al. 2017). The use of a LATER model was largely motivated by its relative simplicity and the ability to quantify the parameters that describe motor recruitment and its association with movement initiation and not necessarily indicative of the underlying neural mechanisms per se. Indeed, recording of the neurons in the primary and premotor cortex, unlike those in the oculomotor system (Dorris et al. 1997; McPeek and Keller 2002a, 2002b; Murthy et al. 2007; Nelson et al. 2016; Thompson et al. 1997), are better understood as emerging from a population of activity distributed across brain areas as part of a dynamic ensemble (Churchland et al. 2010; Georgopoulos et al. 1986; Kaufman et al. 2014). Here we used the LATER model to test whether the model can provide a framework that links movement initiation to the recruitment of motor units in the periphery, where the latter being a lower dimensional signal may be amenable to such an analysis. As expected from the early delay period activity, the correlations between the parameters of the model and reaction times were modest but consistent and significant within and across subjects. However, and not surprisingly, the correlations became much stronger, the close the activity was to movement initiation. To our knowledge, this approach has been used for the first time, to study how accumulation to threshold models (LATER) can be used to assess whether the recruitment of motor activity observed in the periphery is linked in a systematic manner to movement onsets.

By recording from motor activity using high-density surface arrays in the shoulder muscle, we also tested the generality of our findings across tasks. Longer reaction times in both tasks (delayed and immediate task), in the hand and cursor studies was associated by a decrease in slope. However, we also noticed some difference between the recruitment patterns associated with delayed and immediate movements. The delay period in the delay task for hand movement showed modulations which could be used to prepare for the upcoming movements and hence it was not surprising that faster reaction times was associated with an increase in the baseline activity at the time of the GO cue. However, we also observed some differences between the two tasks, with the slope being the significant parameter for explaining RT differences for hand movements but not for cursor movements. Interestingly, onset was the consistent factor that could explain ensuing RT changes for delayed and immediate tasks during cursor movements, despite the same activity level at the time of GO cue. This implies that the parameters of the LATER model were able to capture the nuances in the pattern of putative motor unit activity that contributed towards changes in RT. Where slope was the important parameter for hand studies that could explain variability in RT, onsets played a consistent part in both the task contexts (delayed and immediate movements) for the cursor task. Further, we also found that threshold activity at the time of movement initiation remained the same for delayed and immediate movements across RT as per the predictions of the LATER model, but this value differed depended on whether the condition was a cursor or hand movement. This observation is interesting as task specific affects have also been reported in different brain areas for eye and hand movements (Basu et al. 2021; Mushiake et al. 2009; Schlag-Rey et al. 1997; Tanji et al. 2007).

### Methodological Issues

It is noteworthy and rather striking that the results we obtained have not been documented in prior EMG studies in humans of reaching movements using either surface EMG or even single unit recordings of motor units. We believe several factors may have contributed to our ability to detect these subtle changes that may have gone unnoticed in prior work. First, most of the previous studies have used global EMG measures such as root mean square (rms) from to study recruitment of motor units. These measures are generally crude and summate EMG signal over larger surface area owning to the size of surface electrodes used and thus might not be sensitive enough to detect local or small changes in the muscle response as detected by point contact electrodes within the grid array (as seen in log scale, see inset **Fig 1E**), and thus, evidence of early recruitment may have been missed. Our results suggest that the noise or the variability present in the RMS has certain features which could be exploited by a thresholding operation to isolate point processes that more closely align with the motor unit recruitment. However, an obvious shortfall of such a thresholding approach would be to render the data not sensitive to decreases in motor activity. Perhaps, such a bias would have the fortuitous effect of amplifying any underlying recruitment that would be otherwise obscured by decrements in EMG activity. Second, the use of spatial filtering techniques for high-density surface array EMG recordings is known to selectively enhance local voltage changes which would be more likely to reflect the recruitment patterns of motor units (Gazzoni et al. 2004; Luca et al. 2006; Mambrito and De Luca 1984; Masuda and De Luca 1991; Merletti and Farina 2009; Rau and Disselhorst-Klug 1997) rather than global muscle activity; and our use of template matching and the measurement of the propagation of the signal along the muscle fibres suggests this to be the case. In this context, it is also worth mentioning that our interelectrode spacing of 2.5 mm is much smaller than the typical distances used in many grid arrays and could have contributed to isolating a larger proportion of smaller motor units. Third, unlike many previous reports, our study focussed on proximal shoulder muscles that may be preferentially targeted by the reticulospinal system that may be responsible for the observed delayed activity since these muscles also robustly express a stimulus locked response (Kearsley et al. 2021). Finally, since this early activity was correlated with RT, had spatially selective response fields (see **<u>Fig S5</u>**) and was only observed for delayed movements but not for cursor movements generated by isometric muscle force, it cannot be an artifact.

## Acknowledgement

This study was supported by an Intensification of Research in High Priority Areas Grant from the Department of Science and Technology, Government of India; a Department of Biotechnology - Indian Institute of Science (DBT-IISc) phase 2 partnership programme grant; and institutional support from the Ministry of Human Resource Development to A.M. and S.P.R. were supported by scholarship by Indian Institute of Science, Bangalore. We also thank the anonymous reviewers for their inputs.

## Author Contributions

S.P.R. and A.M. designed the experiments, S.P.R. performed the experiments and analysed the data and both the authors contributed to writing the manuscript.

