## Supplementary figures for "Shoulder muscle recruitment of small amplitude motor units during the delay period encodes a reach movement plan which is sensitive to task context"

(Supplementary figures and legends)

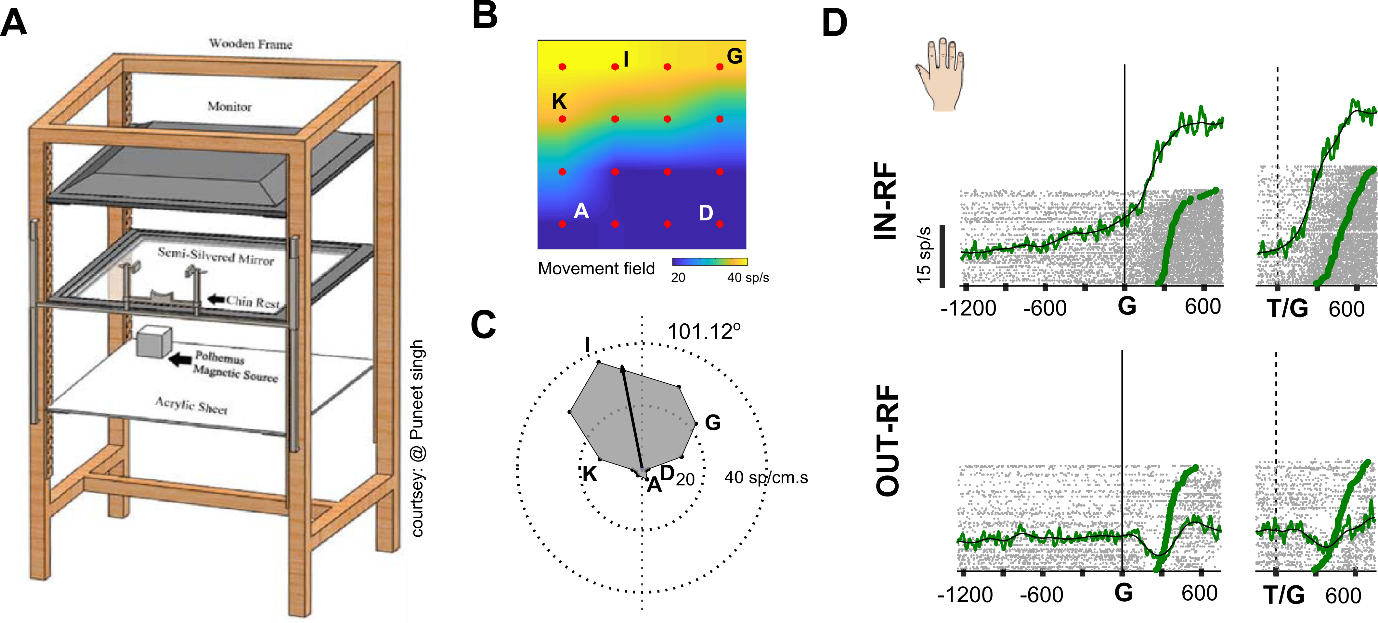

**S1: Reach movement experimental setup and response fields of Anterior Deltoid muscle activity**

A) Schematic representation showing the experimental setup for hand movements. B) Example from one of the subjects showing a typical movement field generated in the workspace when hand movements were made to different target locations arranged on an imaginary rectangular grid. Anterior Deltoid muscle activity is color coded with yellow indicating high activity and blue indicating lower activity. C) A polar plot showing the preferred movement direction. D) Top panel: Response of the muscle when the target was in the movement field for delayed (left panel) and immediate movements (right panel). Bottom panel: Response of the muscle when the target was located away from the movement field.

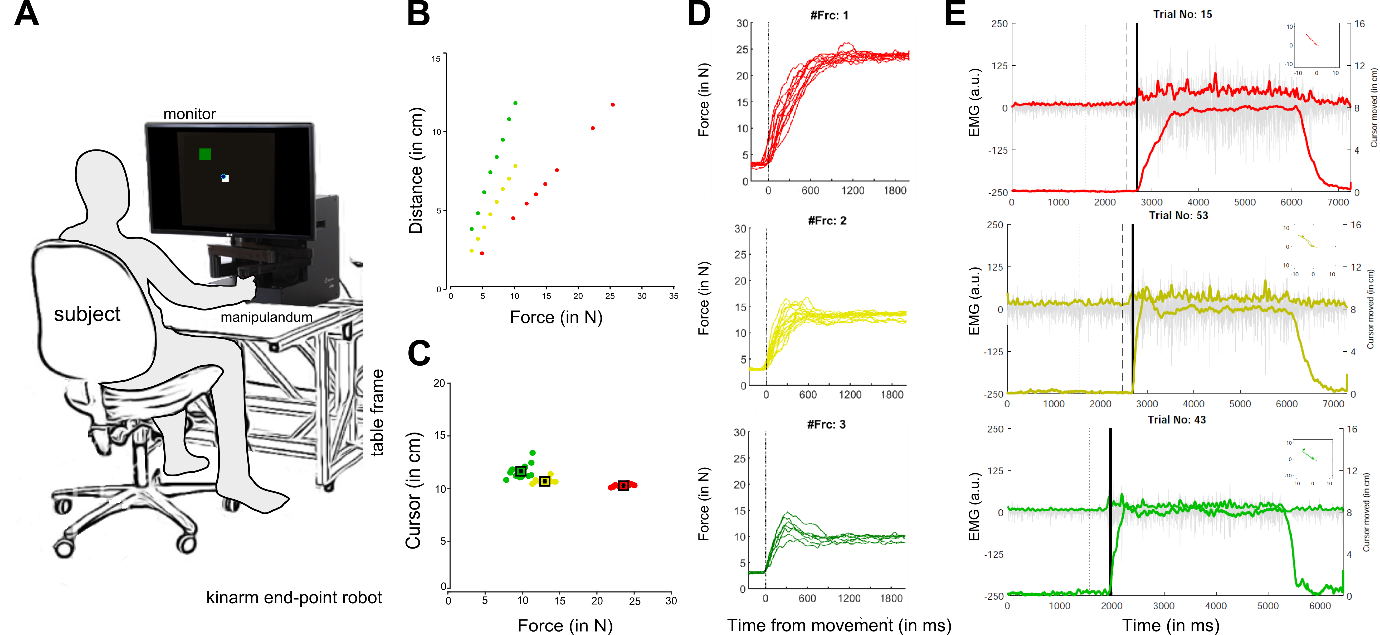

**S2: Isometric cursor movements and EMG activity at different force levels.**

A) Schematic representation showing the experimental setup for cursor movements. B) Scatter plot showing distance moved by the cursor on the screen as a function of different force levels applied on the robotic arm. C) The distance moved by the cursor from the centre of the fixation spot at the end of each trial for each force condition. D) Top panel (red), middle panel (yellow), and bottom panel (green) showing example trials as a function of time, aligned on movement onset for each force level, respectively. E) Root mean square (rms) response variation in the strength of the EMG signal for example trials as a function of different force levels (red, yellow and green).

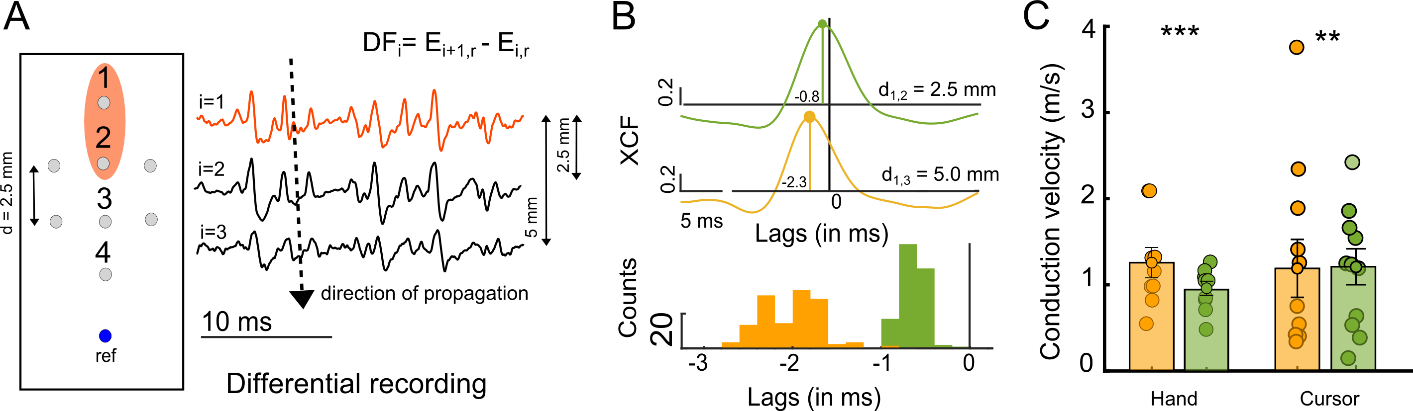

**S3: sEMG array recordings and estimation of conduction velocity**

A) Left panel. A schematic representation of a high-density array used for recording surface electromyographic (emg) signals from the anterior deltoid muscle. The reference electrode was towards the insertion point of the muscle. Middle panel. A snippet showing the propagation of signal along the direction of electrodes (1, 2, 3, and 4) when analysed in differential recording mode. Right panel. Cross-correlation plots showing a lag between signals recorded in differential mode with two electrodes 2.5 mm apart (green) or 5 mm apart (orange). Histogram from an example session, showing the distribution of time lags or conduction delay between electrodes that were separated by 2.5 mm (green) and 5.0 mm (orange). D) Bar plot showing the average conduction velocity estimated between electrodes separated by 2.5 mm (green) and 5.00 mm (orange).

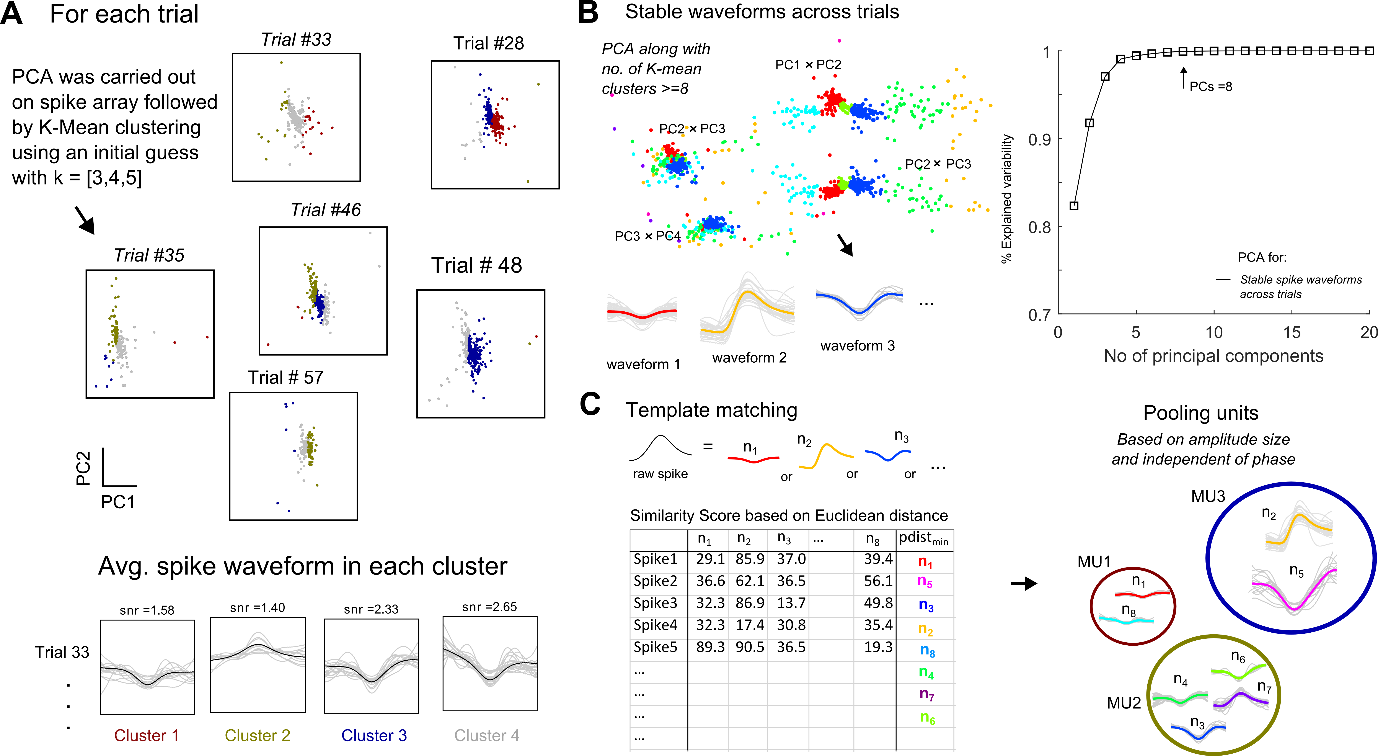
**S4: Spike sorting algorithm and isolation of motor units based on amplitude size**

A) Shows average spike waveforms (k=4) isolated per trial using principal component analysis (PCA) followed by K-means clustering method. Top Panel. Shows spikes marked as individual dots (.), for each trial spread across first two PCs or principal component space grouped into (i) red, (ii) grey, (iii) green, and (iv) blue colour clusters respectively. Bottom Panel. Shows the raw spikes and average shape for the waveform for each cluster and for an example trial as lines in grey and black respectively. B) Shows the stable waveforms identified for the example session by pooling and analysing average shape of spikes (obtained in Step A) from each cluster within each trial, using PCA followed by K-means clustering method. Left panel. Shows eight different clusters or stable waveforms identified for the example session as viewed among first 4 different PCA spaces. Right Panel. Shows the minimum number of principal component axes (PCs) required to explain the variability across average spike waveforms across trial. C) Left panel. Shows a snippet from the table containing a similarity score between raw spikes and stable wave forms isolated across trials using least or minimum Euclidean distance. Right Panel. Shows the different units that were pooled based on their peak amplitude size and independent of their phase into small, medium and large amplitude size motor units shown in red, green and blue colours, respectively.

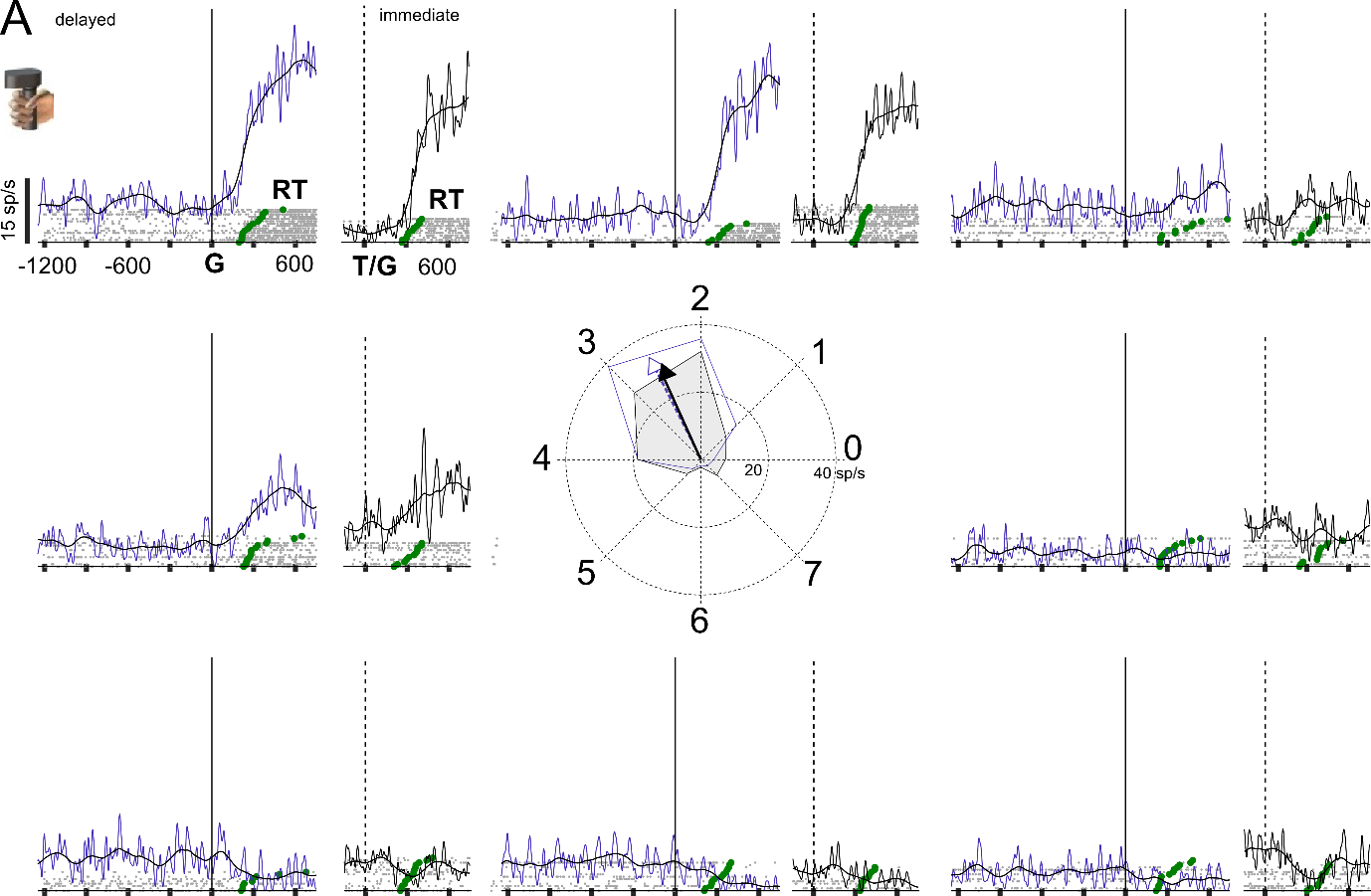

**S5: Spatially tuned response field for cursor-based movements**

A) EMG responses for delayed (blue) and immediate (black) movements, aligned on the go cue, for eight different target locations, for a representative session involving isometric cursor movements. Each grey marker represents a spike. Each spike train represents the response on a single trial which was sorted based on reaction time (green markers). The solid line for the delayed (blue) and immediate task (black) represents the trend seen in the muscle activity for the example session.

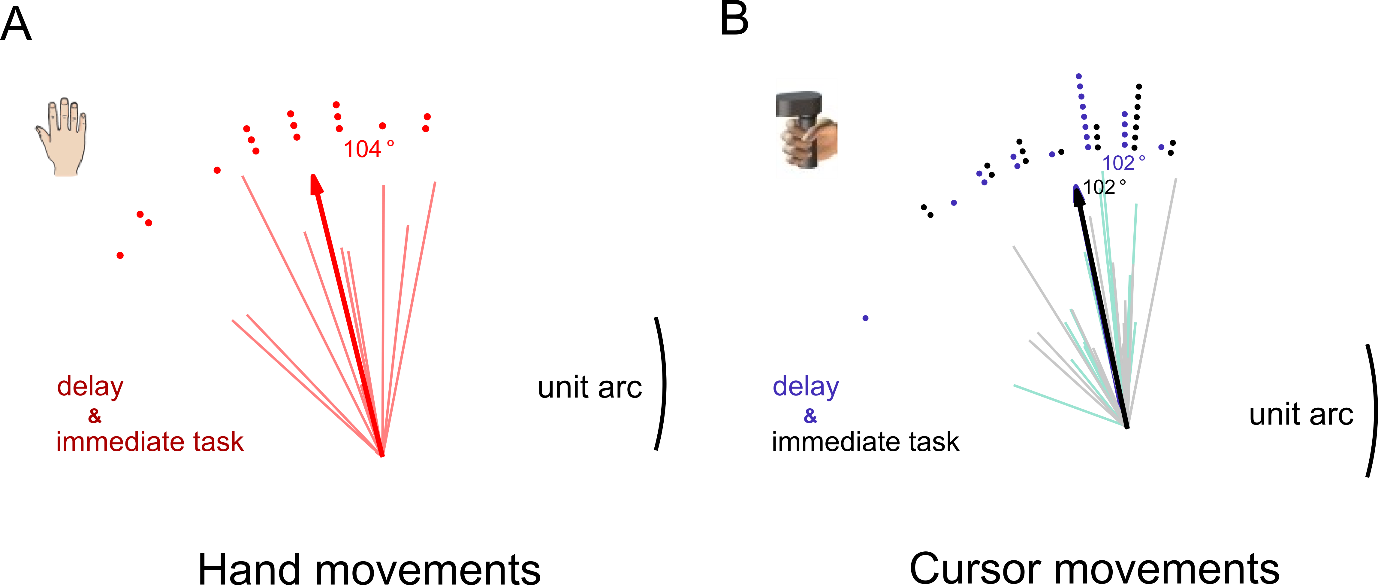

**S6: Preferred movement directions of the anterior deltoid muscle during hand and cursor movements.**

A) The polar plot shows the preferred direction and amplitude of the movement field for the population (thick line) and each session (thin lines) that was recorded prior to delayed and immediate hand movements. The dots demarcate the frequency of occurrence. B) The polar plot shows the preferred direction of the movement field for the population (thick and dashed line) and each session (thin lines) for immediate (grey) and delayed (blue) cursor movements, respectively. The dots demarcate the frequency of occurrence.

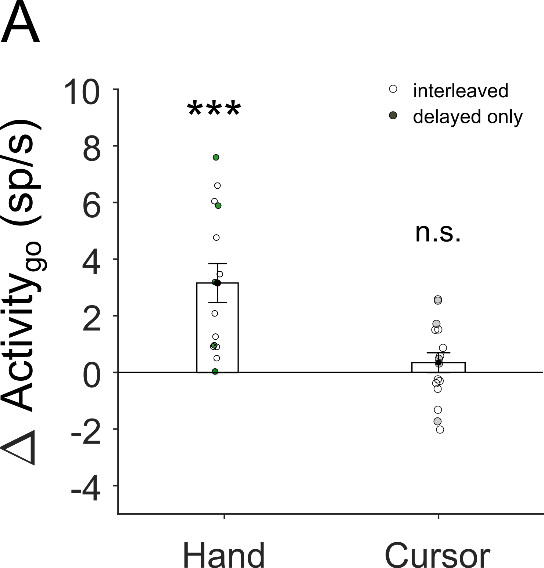

**S7: Changes in motor activity for anterior deltoid muscle during the delay for hand and cursor-based movements.**

A) The bar plot shows the increase and absence of an increase in motor activity during the hold time for the hand (Δ Activity at go: 4.0±0.9, t(13)=4.6, p<0.001) and cursor movements (Δ Activity at go: 0.34±0.35, t(15)=0.98, p=0.34 respectively, with interleaved (unfilled, For hand: 9 subjects and for cursor: 11 subjects) and delayed only task conditions (filled circles, , For hand: 5 subjects and for cursor: 5 subjects). *n.s.* means not significant; * means P<0.05; ** means P<0.01; *** means P<0.001.

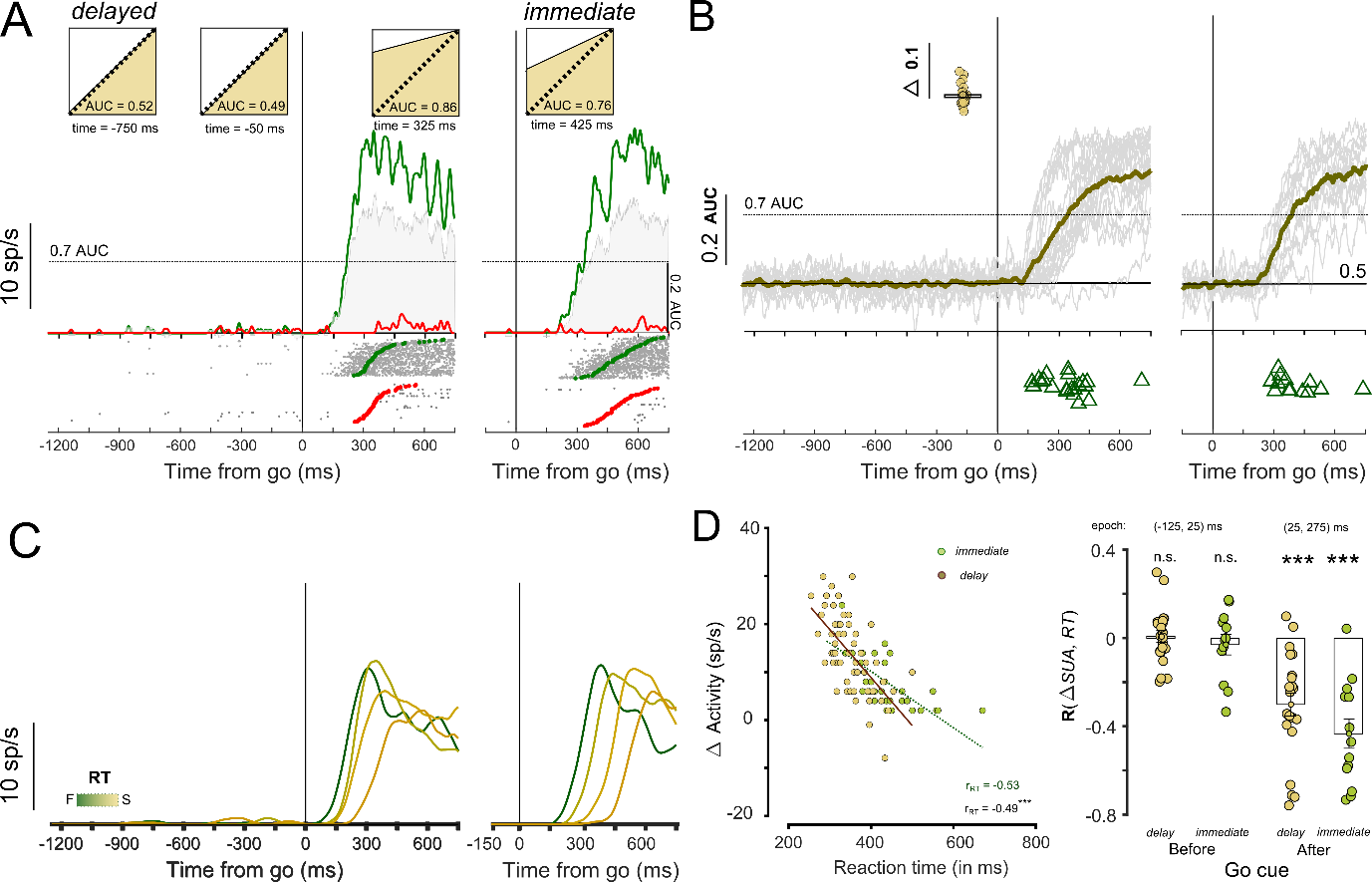

**S8: Spatially selective responses and temporal correlations for non-rampers from reach experiments during the reaction time.**

A) A representative session showing non-ramping motor unit activity for delayed and immediate hand movements, aligned on the go cue for movements made towards (green) and away (red) from the movement field. The inset below shows the spiking activity of the motor unit for each trial marked in grey and the time when the movements were initiated towards (green circles) or away (red circles) from the movement field. The grey shaded area shows the discriminability between in-MF and out-MF response for the EMG signal measured using the area under the curve (AUC) values at each time bin. The inset above shows the separability between in-MF and out-MF response during baseline (-750ms), prior to go cue (-50ms) and following go cue (325ms) for the representative motor unit. B) Shown in grey are the AUC values from the ROC analysis for EMG responses in-MF and out-MF across different sessions for delayed (left) and immediate responses (right). The solid line (brown) shows the average mean value for the population. Inset. Shows the bar plot for the average increase in AUC value from baseline prior to the go cue. Triangles (in olive) mark the onset for direction discriminability for increasing trend during the delay period for AUC values across all sessions. Triangles (in green) show the direction discriminability onsets during the reaction time period based on an AUC>0.7 that demarcates a second later phase of recruitment. C) Activity for the ramping motor unit during delayed and immediate movements, aligned on the go cue for different reaction times (RTs). Inset: Show trial by trial scatter plot (olive) for change in activity from baseline prior to go cue as a function of reaction time. D) Left panel. Scatter plots of ramping activity following the go cue for delayed (olive) and immediate movements (green) as a function of reaction time for the representative session. Right Panel. Bar plot for Pearson’s correlations coefficient between change in activity from baseline prior to the go cue and reaction time, for delayed (olive) and immediate hand movements (green).

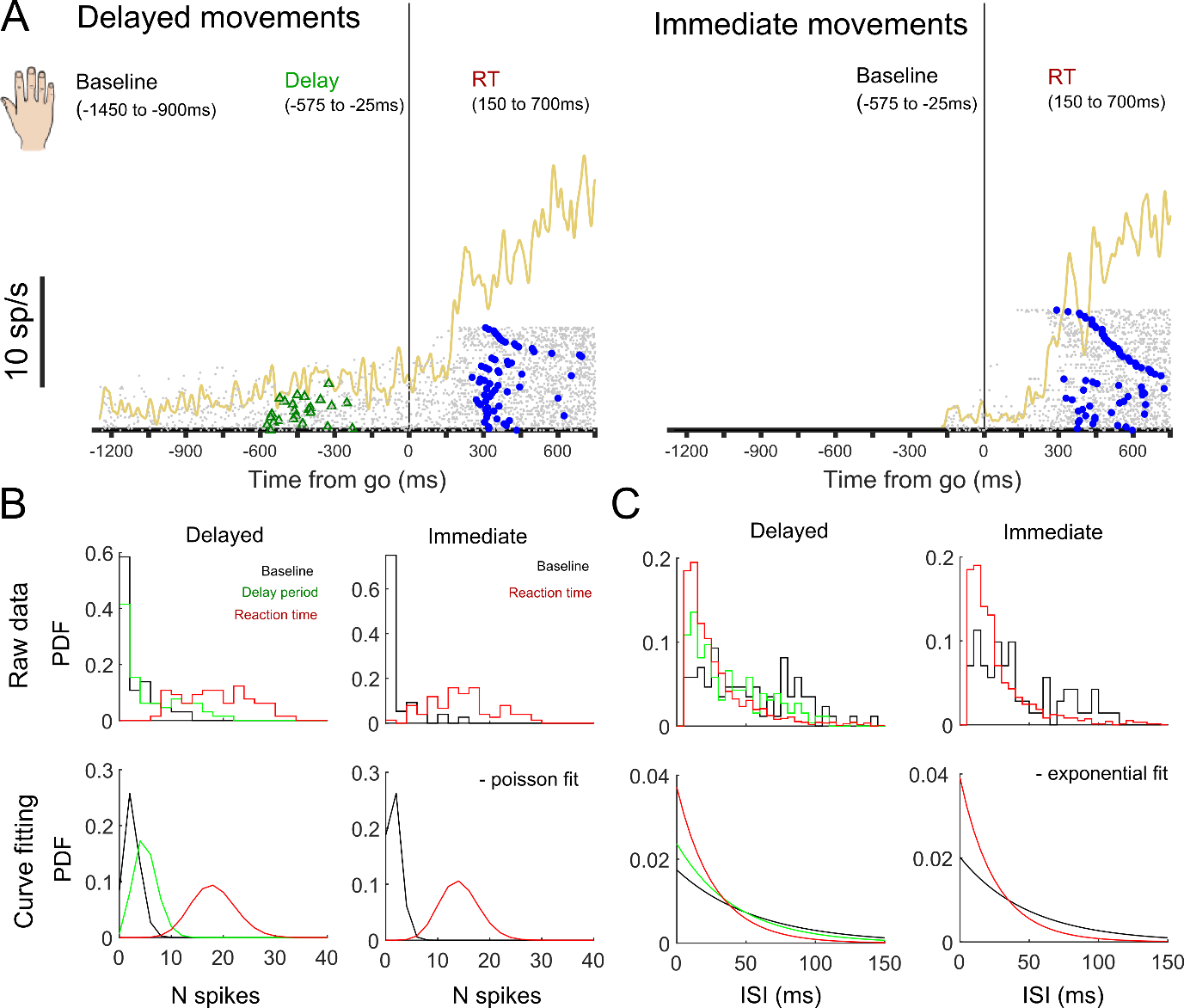

**S9: Poisson analyses to measure single trial onsets for different phases in the task.**

A) Example of a motor unit showing different phases of ramping activity and modulation based on the spike pattern during baseline, delay period, and reaction time for delayed and immediate movements. Gray dots in the raster mark the spike train for each trial sorted based on the increase in spike count observed during the delay period. Green (triangles) mark the EMG onset, estimated using Poisson spike train analysis during the delay period for the delayed condition. The blue (circles) markers show the estimated onset for movements following the go cue for each trial. B) Top Panel. Shows the spike count distribution during baseline (black), delay period (green), and reaction time (red) epochs for delayed and immediate task conditions respectively. Bottom panel. Showing estimated spike count (N spikes) for each epoch following a Poisson like distribution and process. C) Top Panel. Same as B. but for inter-spike intervals during baseline (black), delay time (green) and reaction time (red) epochs for delayed and immediate movements respectively. Bottom Panel. Showing estimated inter-spike interval (ISI) times following an underlying exponential distribution in accordance with Poisson process.

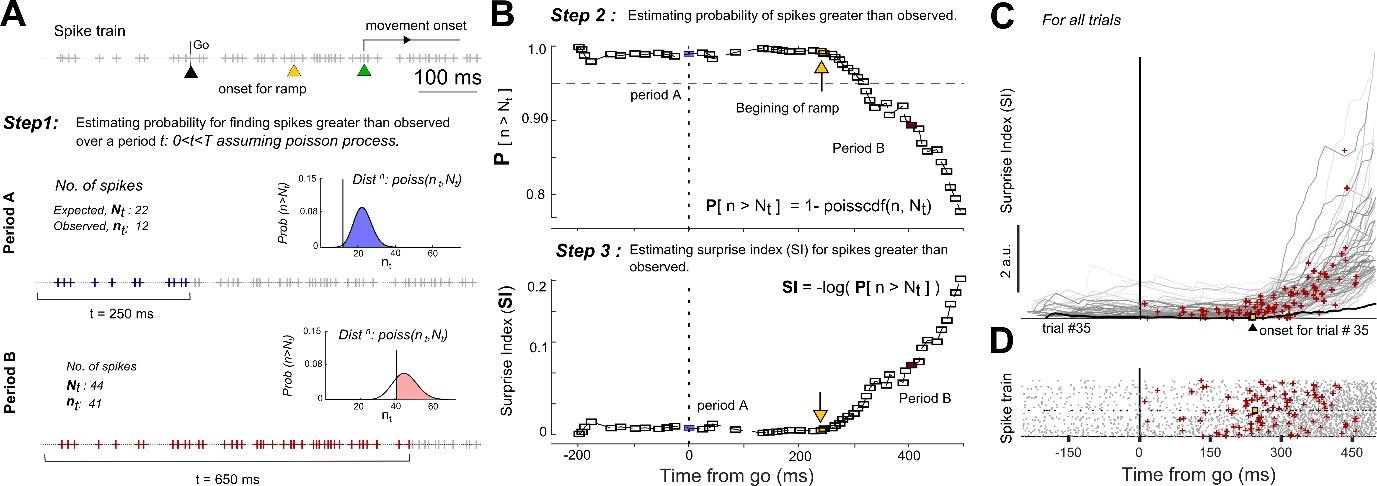

**S10: Poisson spike train algorithm for detecting single trial onsets**

A) Example showing spiking activity of an isolated motor units as a spike train (+, plus sign) marked in gray during the reaction time epoch for a single trial. Triangles in black, yellow, and green mark the go cue, onset for ramping activity as detected by the Poisson spike train onset detection algorithm, and the reaction time for the example trial. Step1 describes the logic for the algorithm used in onset detection for ramping activity for the given spike train. Shown below for period A (in blue, 250ms) and for period B (in red, 650ms) are measures used for estimating average firing rate over the period t: 0<t<T and the probability for finding spikes greater than actually observed during interval t<T. Inset(s) show the spike counts as a Poisson distribution based on the expected and observed number of spikes for the given interval. Blue and red shaded regions estimate the cumulative probability for finding a greater number of spikes than observed for period A and period B, respectively. B) Top Panel. Shows the estimated probability for observing spikes greater than observed as a function of time. Bottom Panel. Shows the surprise Index or negative logarithm of the probability for finding a greater number of spikes than observed as a function of time. The algorithm analyses the local maxima’s and repeats the process in ascending and descending order to mark end and beginning of ramping activity to maximises the SI function to detect and mark the onset time for the ramping activity in spike train. C) The SI across all trials as a function of time and marked onsets (red,+) as detected by the algorithm. The line in black and squares in yellow shows the SI function for the representative trial #35 with the detected onset, respectively. D) Shows the raster plot for each trial showing spikes as grey (dots) and onsets a red (+).

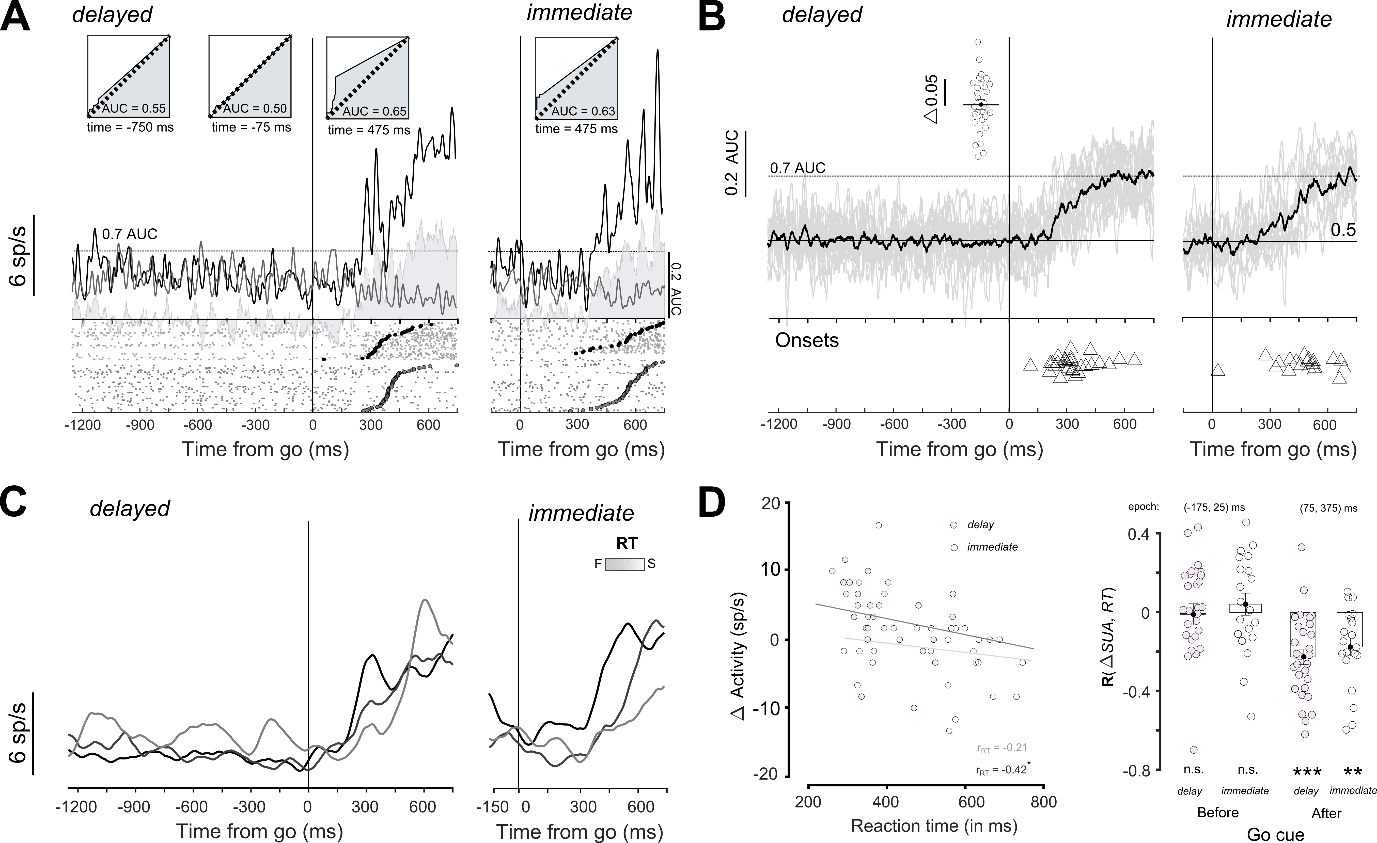

**S11: Spatially selective responses and temporal correlations for non-rampers from the isometric cursor task during reaction time**

A representative session showing non-ramping motor unit activity for delayed and immediate hand movements, aligned on the go cue for movements made towards (black) and away (grey) from the movement field. The format is the same as S8.

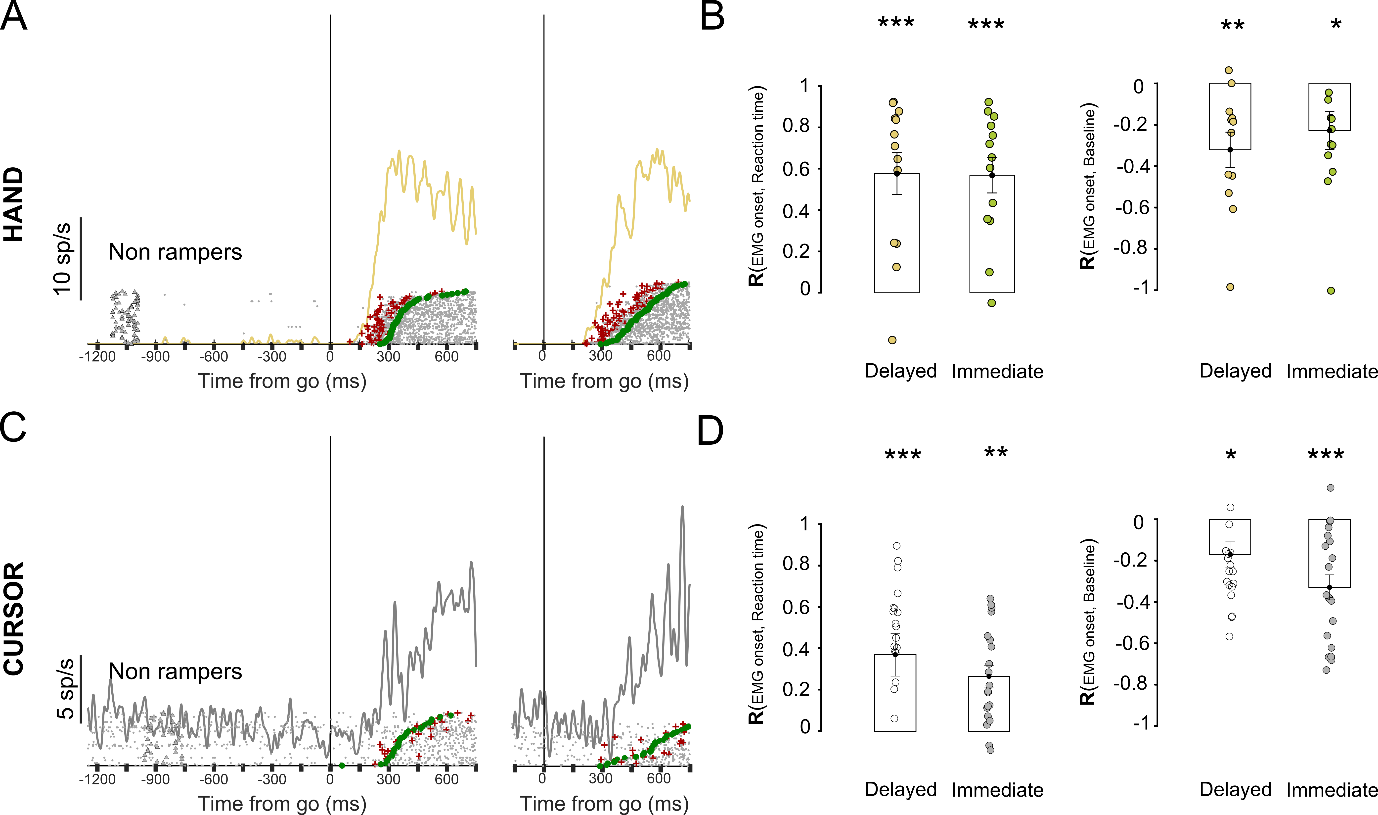

**S12: Non-rampers show correlations between baseline emg activity, onsets in spiking pattern, and time to initiate movements following the go cue.**

A) Example shows the onsets detected for non-ramping motor units during delayed and immediate movements, aligned to the go cue. Grey dots in the raster mark the spike train for each trial sorted based on the increase in spike count observed during the delay period. Red markers (+, plus sign) represent the EMG onset detected using Poisson spike train analysis during the reaction time epoch (following the go cue). The green markers (circles) show the onset of movements following the go cue for each trial. B) Bar plot shows the average Pearson’s correlations coefficient between EMG onsets (following the go cue) estimated using spike train analysis and (i) reaction time (left panel) and (ii) baseline activity (right panel) for delayed (olive) and immediate hand movements (green). C) Example showing EMG onsets detected from a non-ramper from the cursor experiment. D) Same as in B. but for cursor movements. *n.s.* means not significant; * means P<0.05; ** means P<0.01; *** means P<0.001.

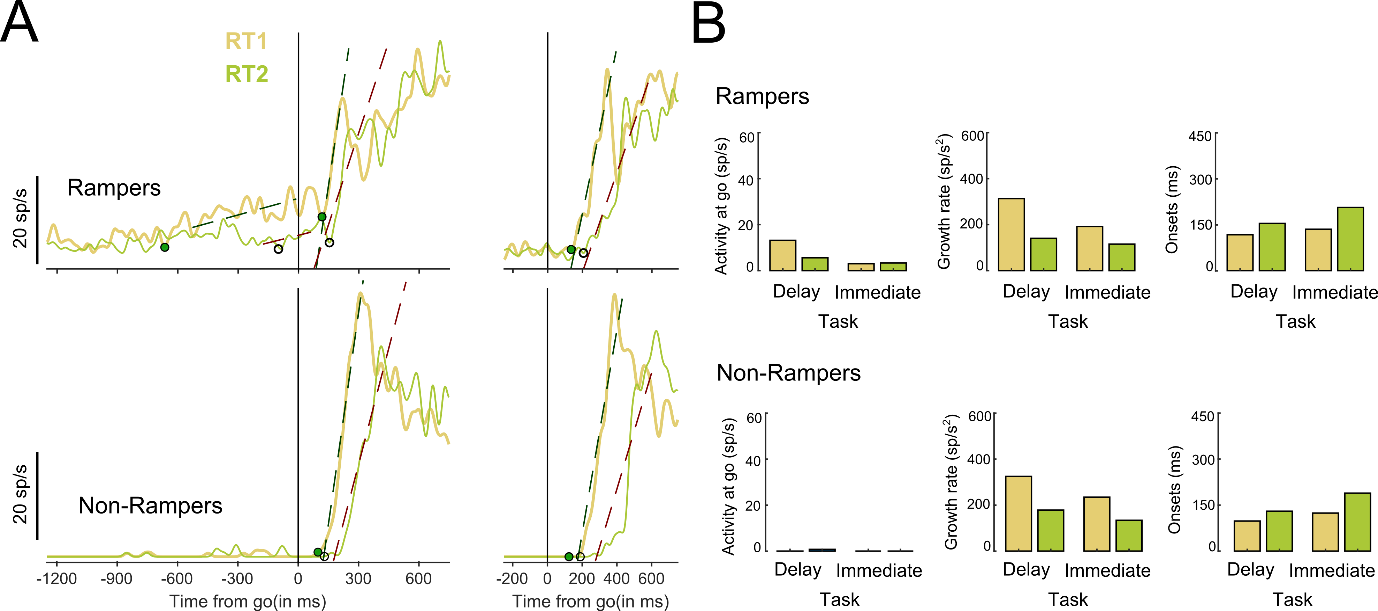

**S13: Baseline activity, changes in slopes and onsets in spiking pattern, as detected by the algorithm to estimate parameters of the accumulator model, to describe motor preparation and initiation of movements prior to and following the go cue.**

A) Top Panel. Example shows the slopes and onsets detected for the ramping motor unit (i) prior to and (ii) following go cue for delayed and immediate movements. Bottom Panel. Example shows the onsets detected for non-ramping motor units during delayed and immediate movements, aligned to the go cue. B) Bar plot shows the Activity at Go cue (first panel), growth rate (second panel) and EMG onsets (third panel) following the go cue estimated for fast (olive) and slow hand movements (green) for rampers (top panel) and non-rampers (bottom panel), respectively.
