## Supplementary table for "Shoulder muscle recruitment of small amplitude motor units during the delay period encodes a reach movement plan which is sensitive to task context"

**Table S1:** Showing mean, standard error (SE) and t-statistics for correlation coefficient between root mean square (RMS) and motor activity (MUA) for anterior deltoid muscle during hand and cursor-based movements respectively.

| *Hand movements* | | | Interleaved task only | | | | | | All sessions | | | | | | | | | | | | |
| --- | --- | --- | --- | --- | --- | --- | --- | --- | --- | --- | --- | --- | --- | --- | --- | --- | --- | --- | --- | --- | --- |
| Variable | | Movements | Obs. | Mean | S.E. | Sample t-test | | | | Obs. | | Mean | | S.E. | | Sample t-test | | | | | |
|  |  |  |  |  |  | t-value | df | p-value | |  | |  | |  | | t-value | | df | | p-value | |
| Correlation coefficient b/w RMS and MUA | | Delayed | 9 | 0.9047 | 0.0419 | 21.59 | 8 | <0.001 | 14 | | 0.9289 | | 0.0281 | | 33.052 | | 13 | | | | <0.001 |
|  |  | Immediate | 9 | 0.9017 | 0.0209 | 43.21 | 8 | <0.001 |  | |  | |  | |  | |  | |  | | |

| *Cursor movements* | | | Interleaved task only | | | | | | | All sessions | | | | | | | | |
| --- | --- | --- | --- | --- | --- | --- | --- | --- | --- | --- | --- | --- | --- | --- | --- | --- | --- | --- |
| Variable | | Movements | Obs. | Mean | S.E. | Sample t-test | | | | Obs. | | Mean | | S.E. | | Sample t-test | | |
|  |  |  |  |  |  | t-value | df | p-value |  | |  | |  | | t-value | | df | p-value |
| Correlation coefficient b/w RMS and MUA | | Delayed | 11 | 0.83 | 0.05 | 16.29 | 10 | <0.001 | 16 | | 0.7998 | | 0.0459 | | 17.4424 | | 15 | <0.001 |
|  |  | Immediate | 11 | 0.70 | 0.04 | 16.43 | 10 | <0.001 |  | |  | |  | |  | |  |  |

**Table S2*.*** *Recruitment of motor units based on amplitude size and changes in firing rate.* *Top panel****:*** *Showing pairwise difference between ramping and non-ramping motor units recorded simultaneously during the same session for hand movements****.*** *Bottom panel****:*** *Showing pairwise difference between different non-ramping motor units recorded simultaneously during the same session for cursor movements****.***

| *Hands movements* | | Interleaved task only | | | | | | | All sessions | | | | | | |
| --- | --- | --- | --- | --- | --- | --- | --- | --- | --- | --- | --- | --- | --- | --- | --- |
| Variable | Motor unit | | Obs. | Mean | S.E. | Paired t-test | | | | Obs. | Mean | S.E. | Paired t-test | | |
|  |  | |  |  |  | t-value | df | p-value | |  |  |  | t-value | df | p-value |
| Amplitude | Ramper | | 9 | 28.45 | 5.54 | -4.60 | 10 | <0.001 | | 13 | 63.01 | 26.45 | -2.41 | 12 | 0.0164 |
| (uV) | Non-ramper | | 9 | 98.39 | 18.06 |  |  |  | | 13 | 235.33 | 93.94 |  |  |  |
| Baseline | Ramper | | 9 | 7.39 | 0.92 | 4.51 | 10 | <0.001 | | 13 | 6.99 | 1.31 | 5.09 | 12 | <0.001 |
| (sp/s) | Non-ramper | | 9 | 1.52 | 0.41 |  |  |  | | 13 | 0.78 | 0.36 |  |  |  |
| Recruitment | Ramper | | 9 | 10.01 | 1.31 | 5.65 | 10 | <0.001 | | 13 | 10.13 | 1.35 | 6.85 | 12 | <0.001 |
| (Δ sp/s) | Non-ramper | | 9 | 1.61 | 0.46 |  |  |  | | 13 | 1.21 | 0.39 |  |  |  |

| *Cursor movements* | | Interleaved task only | | | | | | All sessions | | | | | |
| --- | --- | --- | --- | --- | --- | --- | --- | --- | --- | --- | --- | --- | --- |
| Variable | Motor unit | Obs. | Mean | S.E. | Paired t-test | | | Obs. | Mean | S.E. | Paired t-test | | |
|  |  |  |  |  | t-value | df | p-value |  |  |  | t-value | df | p-value |
| Amplitude | Small | 9 | 21.53 | 3.27 | -3.03 | 8 | 0.008 | 13 | 22.44 | 4.00 | -3.09 | 12 | 0.004 |
| (uV) | Big | 9 | 47.92 | 10.84 |  |  |  | 13 | 51.07 | 12.75 |  |  |  |
| Baseline | Small | 9 | 15.70 | 2.90 | 3.41 | 8 | 0.0046 | 13 | 14.34 | 2.33 | 4.16 | 12 | <0.001 |
| (sp/s) | Big | 9 | 6.90 | 1.71 |  |  |  | 13 | 5.71 | 1.27 |  |  |  |

**Table S3.** *Characterising rampers and non-rampers based on different parameters to measure ongoing changes in motor activity during delay period and reaction time epoch.*

| *Hands movements* | | | | | | Interleaved task | | | | | | | | | | | All sessions | | | | | | | | | | |
| --- | --- | --- | --- | --- | --- | --- | --- | --- | --- | --- | --- | --- | --- | --- | --- | --- | --- | --- | --- | --- | --- | --- | --- | --- | --- | --- | --- |
| Movements | | Period | Parameter | Motor units | | Obs. | | Mean | | S.E. | | two sample t-test | | | | | Obs. | | Mean | | S.E. | | two sample t-test | | | | |
|  | |  |  |  | |  | |  | |  | | t-value | | df | | p-value |  | |  | |  | | t-value | | df | | p-value |
| Delayed | | Before GO cue | Onsets | Ramper | | 9 | | -593.55 | | 45.58 | | -13.02 | | 8 | | <0.001 | 11 | | -546.9 | | 39.993 | | -13.675 | | 10 | | <0.001 |
|  | |  | Growth rate | Ramper | | 9 | | 3.32 | | 0.99 | | 3.35 | | 8 | | 0.01 | 11 | | 4.654 | | 1.22 | | 3.807 | | 10 | | 0.0034 |
|  | |  | Increase in activity | Ramper | | 9 | | 2.786 | | 0.53 | | 5.29 | | 8 | | <0.001 | 11 | | 3.37 | | 0.58 | | 5.784 | | 10 | | <0.001 |
|  | |  |  | Non-ramper | | 13 | | 0.23 | | 0.14 | | 1.59 | | 12 | | 0.137 | 12 | | 0.159 | | 0.1 | | 1.594 | | 21 | | 0.125 |
|  | | After GO cue | Onsets | Ramper | | 9 | | 94.00 | | 28.10 | | 1.71 | | 20 | | 0.05 | 11 | | 98.909 | | 27.417 | | -2.628 | | 31 | | 0.0066 |
|  | |  |  | Non-ramper | | 13 | | 139.53 | | 10.90 | |  | |  | |  | 22 | | 161.181 | | 9.864 | |  | |  | |  |
|  | |  | Activity at go cue | Ramper | | 9 | | 9.18 | | 1.82 | | 4.45 | | 20 | | <0.001 | 11 | | 9.477 | | 1.4908 | | 6.765 | | 31 | | <0.001 |
|  | |  |  | Non-ramper | | 13 | | 1.99 | | 0.48 | |  | |  | |  | 22 | | 1.753 | | 0.334 | |  | |  | |  |
|  | |  | Growth rate | Ramper | | 9 | | -17.18 | | 19.99 | | 4.21 | | 20 | | <0.001 | 11 | | -20.405 | | 16.646 | | -5.275 | | 31 | | <0.001 |
|  | |  |  | Non-ramper | | 13 | | 76.01 | | 12.24 | |  | |  | |  | 22 | | 65.976 | | 8.119 | |  | |  | |  |
|  | |  | Threshold | Ramper | | 9 | | 10.73 | | 2.48 | | -2.16 | | 20 | | 0.042 | 11 | | 10.653 | | 2.256 | | -2.8411 | | 31 | | 0.007 |
|  | |  |  | Non-ramper | | 13 | | 19.97 | | 3.09 | |  | |  | |  | 22 | | 20.058 | | 2.044 | |  | |  | |  |
| Immediate | | After GO cue | Onsets | Ramper | | 9 | | 196.30 | | 22.07 | | -0.48 | | 20 | | 0.63 |  | |  | |  | |  | |  | |  |
|  | |  |  | Non-ramper | | 13 | | 208.15 | | 13.19 | |  | |  | |  |  | |  | |  | |  | |  | |  |
|  | |  | Activity at go cue | Ramper | | 9 | | 6.44 | | 1.51 | | 3.42 | | 20 | | 0.003 |  | |  | |  | |  | |  | |  |
|  | |  |  | Non-ramper | | 13 | | 1.67 | | 0.50 | |  | |  | |  |  | |  | |  | |  | |  | |  |
|  | |  | Growth rate | Ramper | | 9 | | 2.79 | | 18.24 | | -2.69 | | 20 | | 0.014 |  | |  | |  | |  | |  | |  |
|  | |  |  | Non-ramper | | 13 | | 53.76 | | 9.56 | |  | |  | |  |  | |  | |  | |  | |  | |  |
|  | |  | Threshold | Ramper | | 9 | | 11.18 | | 2.55 | | -2.05 | | 20 | | 0.05 |  | |  | |  | |  | |  | |  |
|  | |  |  | Non-ramper | | 13 | | 19.65 | | 2.93 | |  | |  | |  |  | |  | |  | |  | |  | |  |
| *Cursor movements* | | | | | Interleaved task | | | | | | | | | | | | | All sessions | | | | | | | | | |
| Motor unit | Period | | Parameter | Movements | Obs. | | Mean | | S.E. | | two sample t-test | | | | | | | Obs. | | Mean | | S.E. | two sample t-test | | | | |
|  |  | |  |  |  | |  | |  | | t-value | | df | | p-value | | |  | |  | |  | t-value | df | | p-value | |
| Non-ramper | Before GO cue | | Increase in activity | Delayed | 20 | | -0.08 | | 0.28 | | -0.30 | | 19 | | 0.7673 | | | 29 | | -0.06 | | 0.23 | -0.25 | 28 | | 0.803 | |
|  | After GO cue | | Threshold | Delayed | 20 | | 18.42 | | 2.53 | | 1.52 | | 19 | | 0.072 | | | 29 | | 15.87 | | 2.05 | -0.69 | 47 | | 0.753 | |
|  |  | |  | Immediate | 20 | | 17.38 | | 2.52 | |  | |  | |  | | | 20 | | 17.38 | | 2.52 |  |  | |  | |
|  |  | | Growth rate | Delayed | 20 | | 28.97 | | 3.97 | | 2.69 | | 19 | | 0.007 | | | 29 | | 27.09 | | 3.08 | 1.85 | 47 | | 0.035 | |
|  |  | |  | Immediate | 20 | | 18.64 | | 3.19 | |  | |  | |  | | | 20 | | 18.65 | | 3.20 |  |  | |  | |
|  |  | | Onset | Delayed | 20 | | 124.5 | | 17.28 | | -1.9 | | 19 | | 0.036 | | | 29 | | 111.45 | | 14.11 | -2.76 | 47 | | 0.004 | |
|  |  | |  | Immediate | 20 | | 186.1 | | 25.46 | |  | |  | |  | | | 20 | | 186.10 | | 25.47 |  |  | |  | |
|  |  | | Activity at GO cue | Delayed | 20 | | 12.52 | | 2.24 | | -0.26 | | 19 | | 0.599 | | | 29 | | 10.72 | | 1.76 | -0.47 | 47 | | 0.680 | |
|  |  | |  | Immediate | 20 | | 12.67 | | 2.24 | |  | |  | |  | | | 20 | | 12.67 | | 2.24 |  |  | |  | |

**Table S4.** *Showing spatial information encoded in ramping and non-ramping motor units as observed during the delay period and reaction time epoch for hand and cursor-based movements.*

| *Hands movements* | | | | Interleaved task | | | | | | All sessions | | | | | |
| --- | --- | --- | --- | --- | --- | --- | --- | --- | --- | --- | --- | --- | --- | --- | --- |
| Motor unit | Period | Parameter | Movements | Obs. | Mean | S.E. | Paired t-test | | | Obs. | Mean | S.E. | Paired t-test | | |
|  |  |  |  |  |  |  | t-value | df | p-value |  |  |  | t-value | df | p-value |
| RAMPERS | Before GO cue | Onsets | Delayed | 9 | -335.70 | 84.64 | -3.96 | 8 | 0.004 | 11 | -300.00 | 72.60 | -4.13 | 10 | 0.002 |
|  |  | Activity at go cue | Delayed | 9 | 0.06 | 0.01 | 7.39 | 8 | <0.001 | 11 | 0.053 | 0.009 | 5.92 | 10 | <0.001 |
|  | After GO cue | Onsets | Delayed | 9 | 144.00 | 18.15 | -4.61 | 8 | <0.001 | 11 | 125.82 | 39.80 | -2.29 | 18 | 0.02 |
|  |  |  | Immediate | 9 | 242.88 | 28.30 |  |  |  | 9 | 242.88 | 28.30 |  |  |  |
| NON-RAMPERS | After GO cue | Activity at go cue | Delayed | 13 | 0.01 | 0.01 | 1.48 | 12 | 0.1648 | 22 | 0.005 | 0.004 | 1.255 | 21 | 0.223 |
|  |  | Onsets | Delayed | 13 | 270.15 | 23.54 | -5.91 | 12 | <0.001 | 22 | 336.772 | 26.539 | -1.675 | 33 | 0.05 |
|  |  |  | Immediate | 13 | 409.53 | 34.28 |  |  |  | 13 | 409.538 | 34.289 |  |  |  |

| *Cursor movements* | | | | Interleaved task | | | | | | | All sessions | | | | | |
| --- | --- | --- | --- | --- | --- | --- | --- | --- | --- | --- | --- | --- | --- | --- | --- | --- |
| Motor unit | Period | Parameter | Movements | | Obs. | Mean | S.E. | Paired t-test | | | Obs. | Mean | S.E. | pooled t-test | | |
|  |  |  |  | |  |  |  | t-value | df | p-value |  |  |  | t-value | df | p-value |
| NON-RAMPERS | After GO cue | Activity at go cue | Delayed | | 20 | 0.00 | 0.01 | 0.35 | 19 | 0.73 | 29 | 0.0014 | 0.0097 | 0.14 | 28 | 0.88 |
|  |  | Onsets | Delayed | | 20 | 333.85 | 21.27 | -4.41 | 19 | <0.001 | 29 | 335.517 | 21.054 | -3.468 | 47 | <0.001 |
|  |  |  | Immediate | | 20 | 466.90 | 33.95 |  |  |  | 20 | 466.9 | 33.95 |  |  |  |

**Table S5.** Showing s*patial information reaching peripheral musculature earlier for ramping motor units when compared to and non-ramping motor units as observed during the reaction time epoch.*

| *Hands movements* | | | | | Interleaved task | | | | | | All sessions | | | | | |
| --- | --- | --- | --- | --- | --- | --- | --- | --- | --- | --- | --- | --- | --- | --- | --- | --- |
| Movements | Period | Parameter | Motor unit | Obs. | | Mean | S.E. | Pooled t-test | | | Obs. | Mean | S.E. | Pooled t-test | | |
|  |  |  |  |  | |  |  | t-value | df | p-value |  |  |  | t-value | df | p-value |
| Delayed | After GO cue | Onsets | Ramper | 9 | | 144.00 | 18.15 | -3.92 | 20 | <0.001 | 11 | 125.82 | 39.80 | -4.499 | 31 | <0.001 |
|  |  |  | Non-ramper | 13 | | 270.15 | 23.54 |  |  |  | 22 | 336.772 | 26.539 |  |  |  |
| Immediate | After GO cue | Activity at go cue | Ramper | 9 | | 242.88 | 28.30 | -3.499 | 20 | 0.0011 | 9 | 242.88 | 28.30 | -3.499 | 20 | 0.001 |
|  |  |  | Non-ramper | 13 | | 409.53 | 34.28 |  |  |  | 13 | 409.538 | 34.289 |  |  |  |

**Table S6.** *Showing temporal information encoded in ramping and non-ramping motor units as observed during the delay period and reaction time epoch for hand and cursor-based movements.*

| *Hands movements* | | | Interleaved task | | | | | | All sessions | | | | | |
| --- | --- | --- | --- | --- | --- | --- | --- | --- | --- | --- | --- | --- | --- | --- |
| Motor unit | Correlation coefficient b/w Reaction time and | Movements | Obs. | Mean | S.E. | Pooled t-test | | | Obs. | Mean | S.E. | Pooled t-test | | |
|  |  |  |  |  |  | t-value | df | p-value |  |  |  | t-value | df | p-value |
| RAMPERS | Activity at GO cue | Delayed | 9 | -0.20 | 0.03 | -6.52 | 8 | 0.002 | 11 | -0.16 | 0.04 | -4.08 | 10 | 0.00 |
|  |  | Immediate | 9 | -0.01 | 0.07 | -0.10 | 8 | 0.920 |  |  |  |  |  |  |
|  | Reaction time | Delayed | 9 | -0.19 | 0.06 | -3.09 | 8 | 0.014 | 11 | -0.17 | 0.06 | -2.59 | 10 | 0.02 |
|  |  | Immediate | 9 | -0.20 | 0.05 | -4.13 | 8 | 0.003 |  |  |  |  |  |  |
| NON-RAMPERS | Activity at GO cue | Delayed | 13 | -0.05 | 0.12 | -0.46 | 12 | 0.65 | 22 | -0.01 | 0.09 | -0.17 | 21 | 0.87 |
|  |  | Immediate | 13 | -0.08 | 0.13 | -0.64 | 12 | 0.54 |  |  |  |  |  |  |
|  | Reaction time | Delayed | 13 | -0.30 | 0.06 | -4.81 | 12 | <0.001 | 22 | -0.16 | 0.07 | -2.35 | 21 | 0.03 |
|  |  | Immediate | 13 | -0.30 | 0.07 | -4.08 | 12 | 0.002 |  |  |  |  |  |  |

| *Cursor movements* | |  | Interleaved task | | | | | | All sessions | | | | | |
| --- | --- | --- | --- | --- | --- | --- | --- | --- | --- | --- | --- | --- | --- | --- |
| Motor unit | Correlation coefficient b/w Reaction time and | Movements | Obs. | Mean | S.E. | sample t-test | | | Obs. | Mean | S.E. | sample t-test | | |
|  |  |  |  |  |  | t-value | df | p-value |  |  |  | t-value | df | p-value |
| NON-RAMPERS | Activity at GO cue | Delayed | 20 | 0.00 | 0.06 | -0.015 | 19 | 0.98 | 29 | -0.01 | 0.05 | -0.17 | 28 | 0.87 |
|  |  | Immediate | 20 | 0.43 | 0.06 | 0.75 | 19 | 0.45 |  |  |  |  |  |  |
|  | Reaction time | Delayed | 20 | -0.20 | 0.05 | -3.94 | 19 | <0.001 | 29 | -0.22 | 0.04 | -5.66 | 28 | <0.001 |
|  |  | Immediate | 20 | -0.17 | 0.04 | -3.82 | 19 | 0.001 |  |  |  |  |  |  |

**Table S7.** *Showing single trial onsets in ramping and non-ramping motor units as detected using Poisson spike train analysis during the delay period and reaction time epoch for hand and cursor-based movements.*

| *Hands movements* | | | | Interleaved task | | | | | | All sessions | | | | | |
| --- | --- | --- | --- | --- | --- | --- | --- | --- | --- | --- | --- | --- | --- | --- | --- |
| Motor unit | Period | Parameter | Movements | Obs. | Mean | S.E. | Paired t-test | | | Obs. | Mean | S.E. | Paired t-test | | |
|  |  |  |  |  |  |  | t-value | df | p-value |  |  |  | t-value | df | p-value |
| RAMPERS | Before GO cue | Onsets | Delayed | 9 | -25.46 | 25.03 | -12.99 | 8 | <0.001 | 11 | -15.93 | 21.40 | -14.76 | 10 | <0.001 |
|  | After GO cue | Onsets | Delayed | 9 | 171.93 | 20.45 | 8.40 | 8 | <0.001 | 11 | 169.29 | 16.79 | 10.09 | 10 | <0.001 |
|  |  |  | Immediate | 9 | 244.28 | 33.08 | 7.38 | 8 | <0.001 |  |  |  |  |  |  |
| NON-RAMPERS | After GO cue | Onsets | Delayed | 13 | 308.47 | 14.03 | 21.97 | 12 | <0.001 |  |  |  |  |  |  |
|  |  |  | Immediate | 13 | 393.69 | 11.56 | 34.03 | 12 | <0.001 |  |  |  |  |  |  |

| *Cursor movements* | | | | Interleaved task | | | | | | All sessions | | | | | |
| --- | --- | --- | --- | --- | --- | --- | --- | --- | --- | --- | --- | --- | --- | --- | --- |
| Motor unit | Period | Parameter | Movements | Obs. | Mean | S.E. | sample t-test | | | Obs. | Mean | S.E. | sample t-test | | |
|  |  |  |  |  |  |  | t-value | df | p-value |  |  |  | t-value | df | p-value |
| NON-RAMPERS | After GO cue | Onsets | Delayed | 20 | 257.13 | 11.75 | 21.87 | 19 | <0.001 | 29 | 262.26 | 8.92 | 29.39 | 28 | <0.001 |
|  |  |  | Immediate | 20 | 279.76 | 19.73 | 14.17 | 19 | <0.001 | 20 | 279.76 | 19.73 | 14.17 | 19 | <0.001 |

**Table S8.** *Showing correlations between single trial onsets, baseline activity and initiation of movements for ramping and non-ramping motor units as detected using Poisson spike train analysis during the delay period and reaction time epoch for hand and cursor-based movements.*

| *Hands movements* | | | | | Interleaved task | | | | | | | All sessions | | | | | | | | | | | | |
| --- | --- | --- | --- | --- | --- | --- | --- | --- | --- | --- | --- | --- | --- | --- | --- | --- | --- | --- | --- | --- | --- | --- | --- | --- |
| Motor unit | Period | Correlation coefficient b/w EMG onset and | Movements | Obs. | | Mean | S.E. | Paired t-test | | | | Obs. | | Mean | | S.E. | | Paired t-test | | | | | | |
|  |  |  |  |  | |  |  | t-value | df | | p-value | |  | |  | |  | | t-value | | | df | p-value | |
| RAMPERS | Before GO cue | Baseline | Delayed | 9 | | -0.19 | 0.07 | -3.39 | 8 | | 0.01 | | 11 | | -0.216 | | 0.064 | | -3.383 | | | 10 | 0.007 | |
|  |  | Reaction time | Delayed | 9 | | -0.12 | 0.08 | -1.43 | | 8 | 0.20 | 11 | | -0.13 | | 0.08 | | -1.65 | | | 10 | | | 0.13 |
|  | After GO cue | Baseline | Delayed | 9 | | -0.17 | 0.03 | -5.70 | | 8 | <0.001 | 11 | | -0.1765 | | 0.0252 | | -4.11 | | | 10 | | | <0.001 |
|  |  |  | Immediate | 9 | | -0.47 | 0.04 | -1.88 | | 8 | <0.001 |  | |  | |  | |  | | |  | | |  |
|  |  | Reaction time | Delayed | 9 | | 0.51 | 0.07 | 7.18 | | 8 | <0.001 | 11 | | 0.45 | | 0.08 | | 5.93 | | | 10 | | | <0.001 |
|  |  |  | Immediate | 9 | | 0.55 | 0.06 | 9.60 | | 8 | <0.001 |  | |  | |  | |  | | |  | | |  |
| NON-RAMPERS | After GO cue | Baseline | Delayed | 13 | | -0.22 | 0.09 | -2.50 | | 12 | 0.02 | 22 | | -0.31 | | 0.08 | | -4.01 | | | 21 | | | <0.001 |
|  |  |  | Immediate | 13 | | -0.32 | 0.08 | -2.47 | | 12 | 0.03 |  | |  | |  | |  | | |  | | |  |
|  |  | Reaction time | Delayed | 13 | | 0.57 | 0.09 | 6.73 | | 12 | <0.001 | 22 | | 0.42 | | 0.07 | | 5.77 | | | 21 | | | <0.001 |
|  |  |  | Immediate | 13 | | 0.57 | 0.11 | 5.28 | | 12 | <0.001 |  | |  | |  | |  | | |  | | |  |

| *Cursor movements* | | | | Interleaved task | | | | | | All sessions | | | | | |
| --- | --- | --- | --- | --- | --- | --- | --- | --- | --- | --- | --- | --- | --- | --- | --- |
| Motor unit | Period | Correlation coefficient b/w EMG onset and | Movements | Obs. | Mean | S.E. | sample t-test | | | Obs. | Mean | S.E. | sample t-test | | |
|  |  |  |  |  |  |  | t-value | df | p-value |  |  |  | t-value | df | p-value |
| NON-RAMPERS | After GO cue | Baseline | Delayed | 20 | -0.19 | 0.07 | -2.43 | 19 | 0.025 | 29 | -0.17 | 0.07 | -2.28 | 28 | 0.03 |
|  |  |  | Immediate | 20 | -0.33 | 0.05 | -6.48 | 19 | <0.001 | 20 | -0.328 | 0.06 | -5.426 | 28 | <0.001 |
|  |  | Reaction time | Delayed | 20 | 0.37 | 0.10 | 3.58 | 19 | 0.002 | 29 | 0.417 | 0.087 | 4.798 | 28 | <0.001 |
|  |  |  | Immediate | 20 | 0.26 | 0.05 | 5.14 | 19 | <0.001 | 20 | 0.26 | 0.05 | 5.14 | 19 | <0.001 |

**Table S8.** Showing t-statistic results of different parameters from the accumulator model for ramping motor unit activity and initiation of movements.

| *Hands movements* | | | |  | Interleaved task | | | | | | All sessions | | | | | |
| --- | --- | --- | --- | --- | --- | --- | --- | --- | --- | --- | --- | --- | --- | --- | --- | --- |
| Motor unit | Period | Parameter | Movements | condition | Obs. | Mean | S.E. | Paired t-test | | | Obs. | Mean | S.E. | Paired t-test | | |
|  |  |  |  |  |  |  |  | t-value | df | p-value |  |  |  | t-value | df | p-value |
| RAMPERS | Before GO cue | Baseline | Delayed | Fast RT | 9 | 12.60 | 3.20 | 0.12 | 8 | 0.9 | 11 | 12.105 | 2.887 | 0.5003 | 10 | 0.6227 |
|  |  |  |  | Slow RT | 9 | 12.52 | 3.52 |  |  |  | 11 | 11.742 | 3.226 |  |  |  |
|  |  | Growth rate | Delayed | Fast RT | 9 | 7.55 | 2.18 | 2.37 | 8 | 0.02 | 11 | 9.252 | 2.343 | 2.859 | 10 | 0.0085 |
|  |  |  |  | Slow RT | 9 | 2.66 | 1.97 |  |  |  | 11 | 3.6324 | 2.0336 |  |  |  |
|  |  | Onsets | Delayed | Fast RT | 9 | -549.60 | 75.59 | 1.73 | 8 | 0.06 | 11 | -561.272 | 68.941 | -1.118 | 10 | 0.1447 |
|  |  |  |  | Slow RT | 9 | -421.20 | 82.80 |  |  |  | 11 | -445.272 | 81.9 |  |  |  |
|  | After GO cue | Activity at go cue | Delayed | Fast RT | 9 | 20.43 | 3.29 | 2.83 | 8 | 0.02 | 11 | 21.33 | 3.06 | 2.83 | 10 | 0.02 |
|  |  |  |  | Slow RT | 9 | 16.43 | 4.07 |  |  |  | 11 | 16.55 | 3.72 |  |  |  |
|  |  |  | Immediate | Fast RT | 9 | 13.54 | 2.96 | 0.55 | 8 | 0.59 | 11 |  |  |  |  |  |
|  |  |  |  | Slow RT | 9 | 12.85 | 3.23 |  |  |  | 11 |  |  |  |  |  |
|  |  | Onsets | Delayed | Fast RT | 9 | 126.1 | 15.76 | -1.79 | 8 | 0.06 | 11 | 117.18 | 18.35 | -2.60 | 10 | 0.01 |
|  |  |  |  | Slow RT | 9 | 158.7 | 23.8 |  |  |  | 11 | 167.27 | 26.32 |  |  |  |
|  |  |  | Immediate | Fast RT | 9 | 156.3 | 22.14 | -6.38 | 8 | <0.001 | 11 |  |  |  |  |  |
|  |  |  |  | Slow RT | 9 | 274.7 | 30.13 |  |  |  | 11 |  |  |  |  |  |
|  |  | Growth rate | Delayed | Fast RT | 9 | 20.12 | 98.77 | 0.37 | 8 | 0.36 | 11 | -4.26 | 90.53 | 0.23 | 10 | 0.41 |
|  |  |  |  | Slow RT | 9 | 0.54 | 55.7 |  |  |  | 11 | -13.95 | 51.64 |  |  |  |
|  |  |  | Immediate | Fast RT | 9 | -32.9 | 64.5 | 0.59 | 8 | 0.28 | 11 |  |  |  |  |  |
|  |  |  |  | Slow RT | 9 | -44.2 | 57.8 |  |  |  | 11 |  |  |  |  |  |
|  |  | Threshold | Delayed | Fast RT | 9 | 26.4 | 3.61 | -0.32 | 8 | 0.75 | 11 | 27.16 | 4.82 | 0.29 | 10 | 0.78 |
|  |  |  |  | Slow RT | 9 | 27.02 | 4.74 |  |  |  | 11 | 24.82 | 5.01 |  |  |  |
|  |  |  | Immediate | Fast RT | 9 | 26.32 | 3.9 | -1.09 | 8 | 0.3 | 11 |  |  |  |  |  |
|  |  |  |  | Slow RT | 9 | 27.54 | 4.6 |  |  |  | 11 |  |  |  |  |  |

**Table S9.** Showing t-statistic results of different parameters from the accumulator model for non-ramping motor unit activity and initiation of movements.

| *Hands movements* | | | |  | Interleaved task | | | | | | All sessions | | | | | |
| --- | --- | --- | --- | --- | --- | --- | --- | --- | --- | --- | --- | --- | --- | --- | --- | --- |
| Motor unit | Period | Parameter | Movements | condition | Obs. | Mean | S.E. | Paired t-test | | | Obs. | Mean | S.E. | Paired t-test | | |
|  |  |  |  |  |  |  |  | t-value | df | p-value |  |  |  | t-value | df | p-value |
| NON-RAMPERS | After GO cue | Activity at go cue | Delayed | Fast RT | 13 | 4.72 | 1.20 | 1.90 | 12 | 0.08 | 22 | 3.98 | 0.79 | 1.73 | 21 | 0.09 |
|  |  |  |  | Slow RT | 13 | 3.48 | 0.90 |  |  |  | 22 | 3.17 | 0.65 |  |  |  |
|  |  |  | Immediate | Fast RT | 13 | 3.17 | 0.90 | -0.32 | 12 | 0.75 |  |  |  |  |  |  |
|  |  |  |  | Slow RT | 13 | 3.28 | 1.03 |  |  |  |  |  |  |  |  |  |
|  |  | Onsets | Delayed | Fast RT | 13 | 103.84 | 6.43 | -4.11 | 12 | <0.001 | 22 | 124.86 | 8.56 | -4.83 | 21 | 0.006 |
|  |  |  |  | Slow RT | 13 | 160.77 | 16.35 |  |  |  | 22 | 174.86 | 12.52 |  |  |  |
|  |  |  | Immediate | Fast RT | 13 | 163.46 | 13.97 | -3.60 | 12 | 0.00 |  |  |  |  |  |  |
|  |  |  |  | Slow RT | 13 | 212.38 | 11.20 |  |  |  |  |  |  |  |  |  |
|  |  | Growth rate | Delayed | Fast RT | 13 | 203.85 | 39.90 | 2.04 | 12 | 0.03 | 22 | 174.46 | 25.23 | 2.71 | 21 | <0.001 |
|  |  |  |  | Slow RT | 13 | 153.06 | 23.48 |  |  |  | 22 | 133.08 | 16.08 |  |  |  |
|  |  |  | Immediate | Fast RT | 13 | 165.41 | 29.00 | 5.86 | 12 | <0.001 |  |  |  |  |  |  |
|  |  |  |  | Slow RT | 13 | 120.07 | 25.90 |  |  |  |  |  |  |  |  |  |
|  |  | Threshold | Delayed | Fast RT | 13 | 30.38 | 6.30 | 0.30 | 12 | 0.76 | 22 | 30.17 | 3.96 | 0.79 | 21 | 0.43 |
|  |  |  |  | Slow RT | 13 | 30.02 | 6.20 |  |  |  | 22 | 29.47 | 3.95 |  |  |  |
|  |  |  | Immediate | Fast RT | 13 | 29.82 | 6.25 | -0.77 | 12 | 0.45 |  |  |  |  |  |  |
|  |  |  |  | Slow RT | 13 | 30.66 | 6.37 |  |  |  |  |  |  |  |  |  |

| *Cursor movements* | | | |  | | Interleaved task | | | | | | | All sessions | | | | | |
| --- | --- | --- | --- | --- | --- | --- | --- | --- | --- | --- | --- | --- | --- | --- | --- | --- | --- | --- |
| Motor unit | Period | Parameter | Movements | | condition | | Obs. | Mean | S.E. | Paired t-test | | | Obs. | Mean | S.E. | Paired t-test | | |
|  |  |  |  | |  | |  |  |  | t-value | df | p-value |  |  |  | t-value | df | p-value |
| NON-RAMPERS | After GO cue | Activity at go cue | Delayed | | Fast RT | | 20 | 24.43 | 4.73 | -1.23 | 19 | 0.23 | 29 | 21.47 | 4.36 | -0.72 | 28 | 0.47 |
|  |  |  |  | | Slow RT | | 20 | 26.98 | 4.49 |  |  |  | 29 | 22.59 | 4.43 |  |  |  |
|  |  |  | Immediate | | Fast RT | | 20 | 25.41 | 4.71 | -0.49 | 19 | 0.62 |  |  |  |  |  |  |
|  |  |  |  | | Slow RT | | 20 | 25.96 | 4.50 |  |  |  |  |  |  |  |  |  |
|  |  | Onsets | Delayed | | Fast RT | | 20 | 94.50 | 26.33 | -2.46 | 19 | 0.01 | 29 | 89.38 | 23.94 | -2.61 | 28 | 0.01 |
|  |  |  |  | | Slow RT | | 20 | 195.15 | 33.41 |  |  |  | 29 | 170.79 | 31.98 |  |  |  |
|  |  |  | Immediate | | Fast RT | | 20 | 160.35 | 20.25 | -2.45 | 19 | 0.01 |  |  |  |  |  |  |
|  |  |  |  | | Slow RT | | 20 | 244.20 | 31.97 |  |  |  |  |  |  |  |  |  |
|  |  | Growth rate | Delayed | | Fast RT | | 20 | 54.91 | 6.17 | 1.05 | 19 | 0.15 | 29 | 55.35 | 6.19 | 2.21 | 28 | 0.02 |
|  |  |  |  | | Slow RT | | 20 | 44.53 | 7.49 |  |  |  | 29 | 38.41 | 6.76 |  |  |  |
|  |  |  | Immediate | | Fast RT | | 20 | 46.62 | 5.97 | 3.75 | 19 | <0.001 |  |  |  |  |  |  |
|  |  |  |  | | Slow RT | | 20 | 29.10 | 5.77 |  |  |  |  |  |  |  |  |  |
|  |  | Threshold | Delayed | | Fast RT | | 20 | 29.20 | 5.21 | -2.14 | 19 | 0.05 | 29 | 25.41 | 4.77 | -1.54 | 28 | 0.13 |
|  |  |  |  | | Slow RT | | 20 | 34.02 | 4.65 |  |  |  | 29 | 28.07 | 4.79 |  |  |  |
|  |  |  | Immediate | | Fast RT | | 20 | 28.46 | 4.74 | -1.22 | 19 | 0.24 |  |  |  |  |  |  |
|  |  |  |  | | Slow RT | | 20 | 30.07 | 4.83 |  |  |  |  |  |  |  |  |  |
